# Characterization of cephalic and non-cephalic sensory cell types provides insight into joint photo- and mechanoreceptor evolution

**DOI:** 10.1101/2021.01.10.426124

**Authors:** Roger Revilla-i-Domingo, Vinoth Babu Veedin Rajan, Monika Waldherr, Günther Prohaczka, Hugo Musset, Lukas Orel, Elliot Gerrard, Moritz Smolka, Matthias Farlik, Robert J. Lucas, Florian Raible, Kristin Tessmar-Raible

## Abstract

Rhabdomeric Opsins (r-Opsins) are light-sensors in cephalic eye photoreceptors, but also function in additional sensory organs. This has prompted questions on the evolutionary relationship of these cell types, and if ancient r-Opsins cells were non-photosensory. Our profiling of cephalic and non-cephalic r-opsin1-expressing cells of the marine bristleworm *Platynereis dumerilii* reveals shared and distinct features. Non-cephalic cells possess a full set of phototransduction components, but also a mechanosensory signature. We determine that Pdu-r-Opsin1 is a Gαq-coupled blue-light receptor. Profiling of cells from *r-opsin1* mutants versus wild-types, and a comparison under different light conditions reveals that in the non-cephalic cells, light – mediated by r-Opsin1 – adjusts the expression level of a calcium transporter relevant for auditory mechanosensation in vertebrates. We establish a deep learning-based quantitative behavioral analysis for animal trunk movements, and identify a light-and r-Opsin-1-dependent fine-tuning of the worm’s undulatory movements in headless trunks, which are known to require mechanosensory feedback.

Our results suggest an evolutionary concept in which r-Opsins act as ancient, light-dependent modulators of mechanosensation, and suggest that light-independent mechanosensory roles of r-Opsins likely evolved secondarily.

## INTRODUCTION

Opsins, a subgroup of G protein-coupled transmembrane receptors (GPCRs), serve as the main light sensors in animal photoreceptor cells. Rhabdomeric Opsins (r-Opsins) are an ancient class of Opsins particularly widespread among invertebrates, typically expressed in larval photoreceptor cells and cephalic eyes that rely on rhabdomeric photoreceptors [1–3]. The role of r-Opsins in light perception has been best studied in the model of the *Drosophila* eye photoreceptor (EP) cells. Stimulation of a light-sensitive chromophore (retinaldehyde) covalently bound to the Gα_q_-coupled r-Opsin apoprotein initiates an intracellular cascade, consisting of a total of 12 proteins, that leads to an increase in intracellular calcium (reviewed in ref. [4]). Whereas most of our knowledge about the function of r-Opsins in animal photoreception stems from studies on cephalic EPs, non-cephalic *r-opsin*- expressing cells are found in representatives of various animal groups. For instance, *r-opsin* homologs demarcate putative photoreceptor cells at the tube feet of sea urchins [5, 6], and in Joseph cells and photoreceptors of the dorsal ocelli of the basal chordate amphioxus [7]. In the case of the brittle star, such non-cephalic photoreceptor cells have been implicated in a form of vision [8]. Yet the diverse locations of *r-opsin*-positive cells, and the fact that they are not strictly associated with pigment cells, have raised the question whether *r-opsin*-positive cells outside the eye might have different functional roles.

This implies the evolutionary question to which extent non-cephalic *r-opsin*- positive cells share an evolutionary history with cephalic eye photoreceptors, or represent independent evolutionary inventions. A biological context in which this question is particularly interesting to address are animals that exhibit segmented body axes, featuring sensory organs in some or all of these segments. Analyses of the early Cambrian Lobopodian fossil *Microdictyon sinicum* suggested that this putative ancestor of arthropods possessed compound eye structures above each pair of legs [9, 10]. This is in line with the idea that segmental photoreceptive organs could have been an ancestral feature, which might have been secondarily modified to allow for a more efficient division of labor between head and trunk. A similar hypothesis could be drawn for ancestors of annelids, a segmented clade of lophotrochozoans: Various recent annelid groups, including opheliids, sabellids and syllids, feature segmental eye spots with rhabdomeric photoreceptors (reviewed in ref. [11]). This would be consistent with the ancient presence of r-opsins and photoreceptive organs in a homonomously segmented annelid ancestor. Given the possible ancestry of segmentation in bilaterians [12–14], the outlined scenarios of segmental photoreceptive organs might even date back to the dawn of bilaterian animal evolution.

However, *r-opsin* genes have also started to be implied in functions that are unrelated to photoreception. Most notably, *r-opsin* genes are expressed in certain classes of mechanosensory cell types, such as the Johnston organ (JO) neurons and the larval Chordotonal organ (ChO) of *Drosophila* [15, 16], or the neuromasts of the lateral line of zebrafish and frog [17, 18]. Experiments assessing functionality of mechanosensation in both JO and ChO neurons have revealed that several *r-opsins* expressed in these receptors are required for proper mechanosensation, and suggest that this function is light-independent [15, 16]. These functional findings add new perspectives to older observations that a subset of mechanosensory cells (to which JO and ChO cells belong) exhibit significant similarities in their molecular specification cascade with eye photoreceptors, comprising analogous use of Pax, Atonal, or Pou4f3 transcription factors [19]. If r-Opsins are to be considered as part of a shared molecular signature in photosensory and mechanosensory cells, this raises divergent possibilities for the evolution of r-Opsin-positive sensory cells: (i) Could r-Opsins have evolved as ancient “protosensory” molecules that were primarily engaged in mechanosensation, only to secondarily evolve to become light receptive, and helping photoreceptors to emerge as a distinct cell type? (ii) Conversely, does the canonical function of r-Opsins in light reception reflect their ancestral role, with r-Opsin-dependent mechanoreceptors representing a secondary evolutionary modification? Or (iii) are there ways in which photosensory functions of r-Opsins could have played an ancient role in mechanoreceptors, even if this role might not be present any more in the investigated *Drosophila* mechanoreceptors?

In order to gain insight into these questions of r-Opsin function, and into the evolution of sensory systems from an independent branch of animal evolution, we characterized *r-opsin*-expressing cells in a lophotrochozoan model system, the marine annelid *Platynereis dumerilii* that is amenable to functional genetic analyses [20–22]. After its pelagic larval stage, *P. dumerilii* inhabits benthic zones [23] that are characterized by a complex light environment, making it likely that light-sensory systems have been evolutionarily preserved in this model, rather than being secondarily reduced. In line with this, *Platynereis* has retained an evolutionarily representative set of *r-opsins* [2, 24] and other photoreceptor genes. The *Platynereis r-opsin1* gene is not only expressed in EPs [2], but also in peripheral cells along the trunk of the animal [17] (cells referred to in this study as trunk *r-opsin1* expressing / TRE cells), making the worm an attractive species for a comparative assessment of r-Opsin function between cephalic and non-cephalic cell types. While both EP and TRE cells express the *gaq* gene that encodes a G_αq_ subunit [17], it has remained elusive whether the TRE cells represent a segmental repetition of the EP cell type along the body plan or represent a distinct sensory modality. Likewise, it is unclear whether r-Opsin1 in TRE cells has a light-sensory role, as in the EPs, or serves a light-independent function, as has been suggested for *Drosophila* JO or ChO neurons.

Here, we established a dissociation and fluorescence-activated cell sorting (FACS) protocol for the *Platynereis* pMos{rops::egfp}^vbci2^ strain that expresses enhanced GFP (EGFP) under the regulatory control of the *Platynereis r-opsin1* gene in both EP and TRE cells [17], and combined this strategy with the targeted mutagenesis of the endogenous *r-opsin1* locus, and an experimental characterization of the r-Opsin1 action spectrum. This allowed us to determine molecular profiles for both EP and TRE cells, revealed that TRE cells, but not EP cells, possess a mechanosensory signature and uncovered that, specifically in the TREs, light – mediated by r-Opsin1 – adjusts the expression level of a plasma membrane calcium transporter relevant for auditory mechanosensation in vertebrates. Our data, therefore, suggest that TRE cells represent a distinct mechanoreceptive cell type, in which r-Opsin1, in difference to the current *Drosophila*-based paradigms, elicits light-dependent functional changes. In line with this, a newly established deep-learning-based approach revealed light-dependent behavioral differences between wildtype and *r-opsin1* mutant trunks. Our results are consistent with the idea that photo- and mechanosensory systems have a common evolutionary origin in a multimodal sensory cell type.

## RESULTS

### Shared and distinct molecular signatures of eye photoreceptor and trunk *r-opsin1*-expressing cells

In order to gain insights into the molecular signatures of EP and TRE cells, we established a mechanical dissociation protocol compatible with FACS, and benchmarked to minimize cell death. We next dissected heads and trunks of the same pMos{rops::egfp}^vbci2^ individuals (**Fig. 1A**), isolated EGFP-positive cells from heads and trunks, and established transcriptomes for both sorted and unsorted cells using Illumina HiSeq sequencing on cDNA amplified by the Smart-Seq2 protocol [25] (**Fig. 1B)**. Gates for FACS (**Fig. 1C,D**) were calibrated using dissociated cells from wildtype heads (**Fig.1-figure supplement 1A**) and trunks (**Fig.1-figure supplement 1B**), to exclude isolation of autofluorescent cells.

**Fig. 1.**
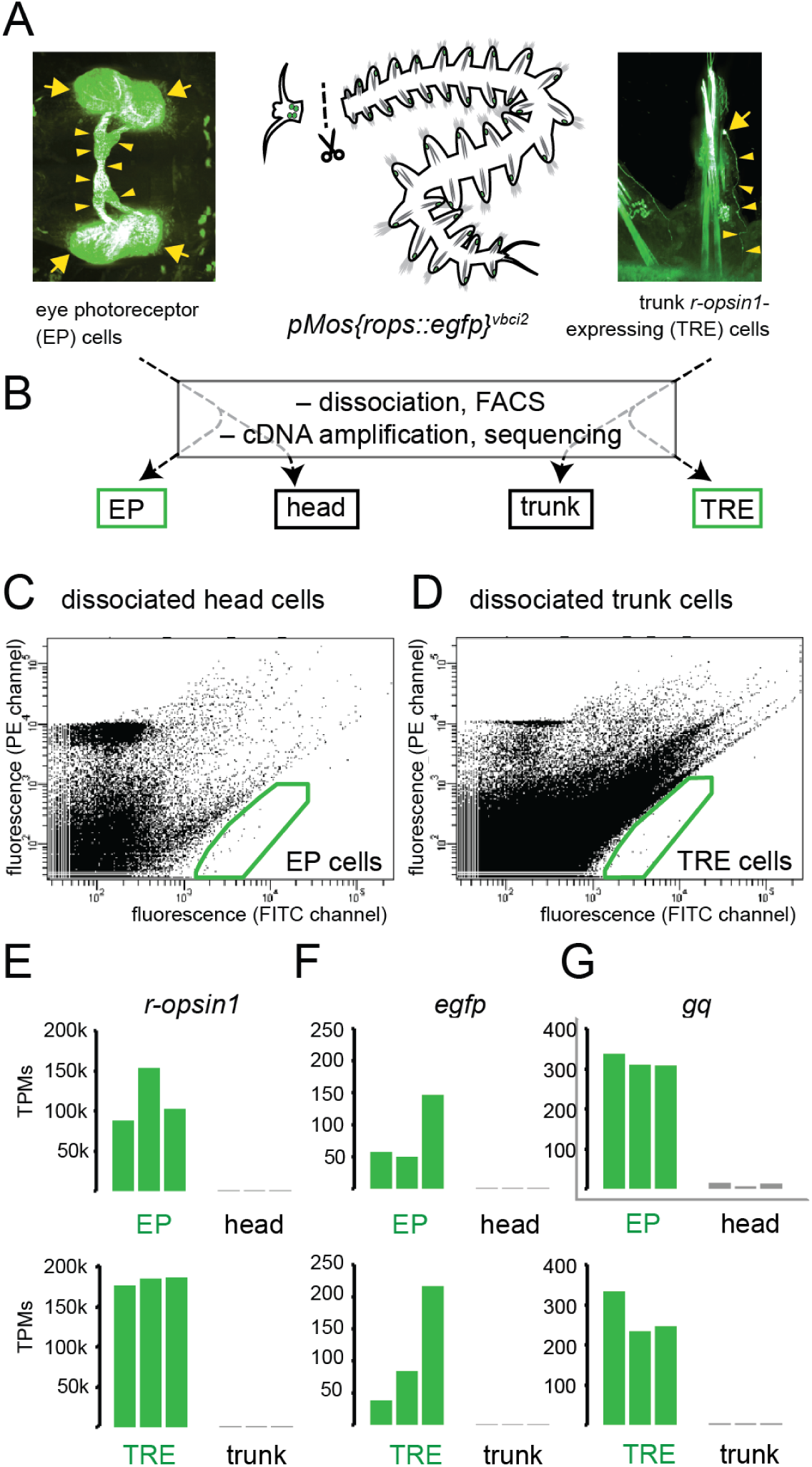
Establishment of molecular signatures of eye photoreceptors and trunk *r-opsin1*-expressing cells. **(A)** Dissection of pMos{rops::egfp}^vbci2^ individuals, separating the head containing eye photoreceptor (EP) cells (left panel) from trunk containing trunk r*-opsin1*-expressing (TRE) cells (right panel). **(B)** Overview over processing and derived cDNA libraries, resulting in FACS-enriched (EP, TRE) and unsorted (head, trunk) samples. **(C,D)** Representative FACS plots showing gated populations (green boxes) of EP and TRE cells, respectively. For non-transgenic controls see **Fig. 1-figure supplement 1A,B**. **(E-G)** comparison of transcripts per million reads (TPM) for the genes *r-opsin1* (E), *enhanced green fluorescent protein/egfp* (F), and *gαq/gq* (G) in individual replicates of EP, head, TRE, and trunk libraries. For comparison of TPMs for non-enriched control genes see **Fig.1-figure supplement 1C,D**. Arrows and arrowheads in (A) designate EGFP-positive cell bodies and projections, respectively.

To validate the sampling strategy, we investigated if this procedure reproduced expected results for genes known to be enriched in both EP and TRE cells. Both *r-opsin1* and *egfp* were up to several thousand times more abundant in libraries derived from EGFP-positive cells than in those of unsorted cells (**Fig. 1E,F**). In further support of successful enrichment, signatures of EGFP-positive cells were consistently enriched in the *gq* gene encoding the G alpha subunit *Gαq* (**Fig. 1G**). *Gq* was previously shown to be strongly expressed in EP and TRE cells [17]. By contrast, the genes encoding the ribosomal subunit Rps9 or the Polo-like Kinase Cdc5, previously established as internal controls for gene expression quantification experiments [26], were not enriched in either EP or TRE cell populations (**Fig. 1-figure supplement 1C,D**).

As these results indicated that the experimental procedure allowed for significant enrichment and profiling of EP and TRE cells, we next used EdgeR [27] to systematically calculate enrichment scores for each of the EGFP- positive populations, compared to the combined set of head and trunk unsorted cells. From a total of 39575 genes, we determined a set of 278 genes (0.7%) to be significantly enriched in EP cells, and a set of 361 genes (0.9 %) significantly enriched in TRE cells (FDR < 0.05) (**Fig. 2A**). 133 genes (0.3 % of total) were shared between the EP and TRE cells (common EP-/TRE-enriched genes), including, expectedly, *r-opsin1* and *gq* (**Figure 2-figure supplement 2**), and leaving 145 (0.4 % of total) EP-specific genes, and 228 (0.6 % of total) TRE-specific genes (**Fig. 2A**). Experiments on selected genes (see Methods: “Analysis and validation of differentially expressed genes”, **Figure 2-figure supplement 1, Figure 2-figure supplement 2**) allowed us to validate the specificity of the predicted sets (**Figure 2-figure supplement 3, Figure 2-figure supplement 4**), pointing at both shared and distinct properties of the *r-opsin1*-expressing cells of the head and the trunk.

**Fig. 2.**
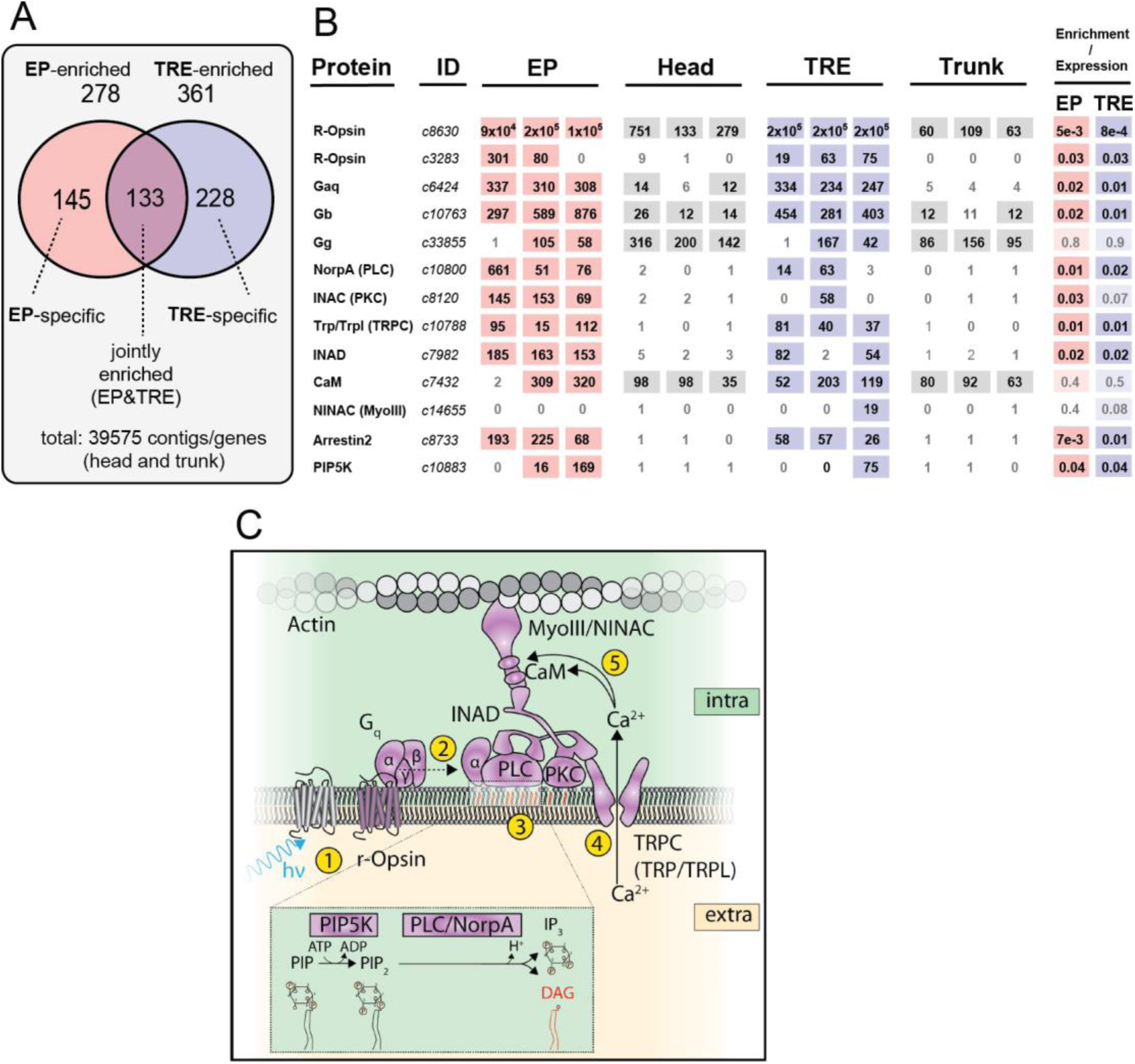
Trunk *r-opsin1*-expressing cells (TRE) share critical elements of the phototransduction cascade with eye photoreceptors (EP). **(A)** Summary of gene/contig counts enriched in EP and TRE cells compared to the combined background (head & trunk; 39575 contigs), and their respective overlap. For expression levels of genes selected for validation see **Fig.2-figure supplement 1**. For validation experiments see Methods: “Analysis and validation of differentially expressed genes”, **Fig.2-figure supplement 2,3,4**. **(B)** Expression and enrichment of phototransduction components in EP and TRE cells. Protein: Key components of the *D. melanogaster* phototransduction pathway (cf. panel C). ID: Corresponding gene ID of the *P. dumerilii* transcriptome. EP, Head, TRE, Trunk: Expression levels (in TPMs) of the respective gene in individual replicates of *P. dumerilii* EP, head, TRE or trunk libraries, respectively. Light shades indicate expression above the established threshold. Enrichment/Expression: FDR values obtained from the differential expression analysis for EP or TRE cells. Dark shades indicate significant enrichment, light shades expression without significant enrichment in the respective cell type. Note that although the *P. dumerilii* best Blast hit to NINAC/MyoIII (*c14655*, E value: 3e-166) is not expressed in EP cells, the second best Blast hit (*c8565*, E value: 1e-64) is expressed in these cells. For sequence identifiers of the relevant *P. dumerilii* genes see **Fig.2-figure supplement 5**. **(C)** Scheme highlighting factors present or enriched in the joint EP/TRE signature (cf. panel B), and their function in critical steps (yellow circles 1-5) of the canonical r-Opsin phototransduction cascade. Enlarged inset shows relevant enzymatic steps in the intracellular leaflet. (C) modelled after ref. [4].

Previous analyses had already revealed several genes to be expressed in adult EPs: *vesicular glutamate transporter (vglut)* [28, 29], the rhabdomeric opsin gene *r-opsin3* [24], the G_o_-type opsin gene *G_o_-opsin1* [21, 30], and the light-receptive cryptochrome gene *l-cry* [26]. *r-opsin3*, *G_o_-opsin1* and *l-cry* were all significantly enriched in the EP-derived transcriptome, when compared to unsorted head cells. While *r-opsin3* has already been described to be expressed in TREs [24], also *G_o_-opsin1*and *l-cry* were part of the specific TRE transcriptome, pointing at an unexpected complexity of light receptors in the TRE cells, and a similar equipment of EP and TRE cells with photoreceptive molecules. Our sequencing data did not cover the published *vglut* gene in any of the samples (possibly reflecting low expression in the adult).

As to potential differences between EPs and TREs, prior analyses had pointed to the expression of circadian clock genes in the EPs and the adjacent brain lobes [26], and both classical and molecular studies suggested the retina as a site of continuous neurogenic activity [31, 32], contrasting with the appearance of the TREs as sparse, differentiated neurons [17]. In line with these expectations, we found the EP-, but not TRE-derived transcriptomes to be enriched, respectively, in the circadian clock gene *bmal*, as well as a homolog of the *embryonic lethal, abnormal vision/elav* gene, a marker characteristic for committed neurons [33, 34].

### EP and TRE cells share a complete phototransduction pathway

Building on these initial results, we next explored if the identified gene sets could provide additional insights into the function and evolution of the TRE cells. We first assessed whether molecular data in addition to the identified photoreceptor molecules would support a possible function of TRE cells in light sensitivity, as it would be expected if these cells represented segmentally repeated cell type homologs of the *P. dumerilii* and *Drosophila melanogaster* EP cells. To test this hypothesis, we compared EP- and TRE-enriched genes of our bristleworm with a published set of genes enriched in *Drosophila* EP cells [35].

Using BLAST-based homology relationships between *D. melanogaster* and *P. dumerilii* genes (see Methods), we established a set of 408 *bona fide P. dumerilii* homologs of the 743 *D. melanogaster* EP-enriched genes. 9 of these were common EP-/TRE-enriched genes. A statistical analysis, based on the generation of 10^4^ sets of 743 randomly-picked *D. melanogaster* genes (see Methods), indicated that this number of common EP-/TRE-enriched genes significantly exceeds random expectation (**Fig. 3-figure supplement 1A**, p = 0.024). Among these 9 overlapping genes, we found 5 *bona fide P. dumerilii* homologs of genes known to be involved in the r-Opsin phototransduction pathway described for Drosophila EP cells [4] (yellow box in **Fig. 3-figure supplement 1B**). By extending our assessment to *bona fide* homologs of additional components of the *Drosophila* r-Opsin phototransduction pathway, we found that putative homologs of 9 and 8 of the 12 components of the r-Opsin phototransduction pathway are enriched in the *P. dumerilii* EP and TRE cells, respectively (**Fig. 2B**). Statistical analysis with 10^4^ random gene sets of matching size (see Methods) revealed these results to be highly significant (p < 10^-4^, for both EP and TRE). Of note, all 12 components of the r-Opsin phototransduction pathway were found to be expressed in the TRE cells of *P. dumerilii* (**Fig. 2B,C**) (p < 10^-4^).

### TRE cells combine photo- and mechanosensory molecular signatures

Following the same strategy, to further explore potential additional functions of the TRE cells, we next tested the molecular relationship between the worm’s TREs and the r-Opsin-expressing, mechanosensory JO neurons of *Drosophila*. For this, we took advantage of 101 genes identified as JO neuron specific in a microarray analysis [15], and 80 *P. dumerilii* homologs of these. Significant subsets of these were found in the common EP-/TRE-enriched signature (9 genes; p < 10^-4^), and the TRE-specific signature (7 genes; p = 1.3×10^-3^) (**Fig. 3A**). The common EP-/TRE-enriched genes essentially reflect the *P. dumerilii* homologs of the aforementioned phototransduction pathway (*rh3/rh4, rh5/rh6, trp/trpl, norpa, gβ76c, pip5k59b*, *arr2* and *klp68D;* **Fig. 3B**). This finding indicates that not only rhodopsin genes are present in JO neurons, as reported [15], but that JO neurons possess a complete r-Opsin phototransduction machinery.

**Fig. 3.**
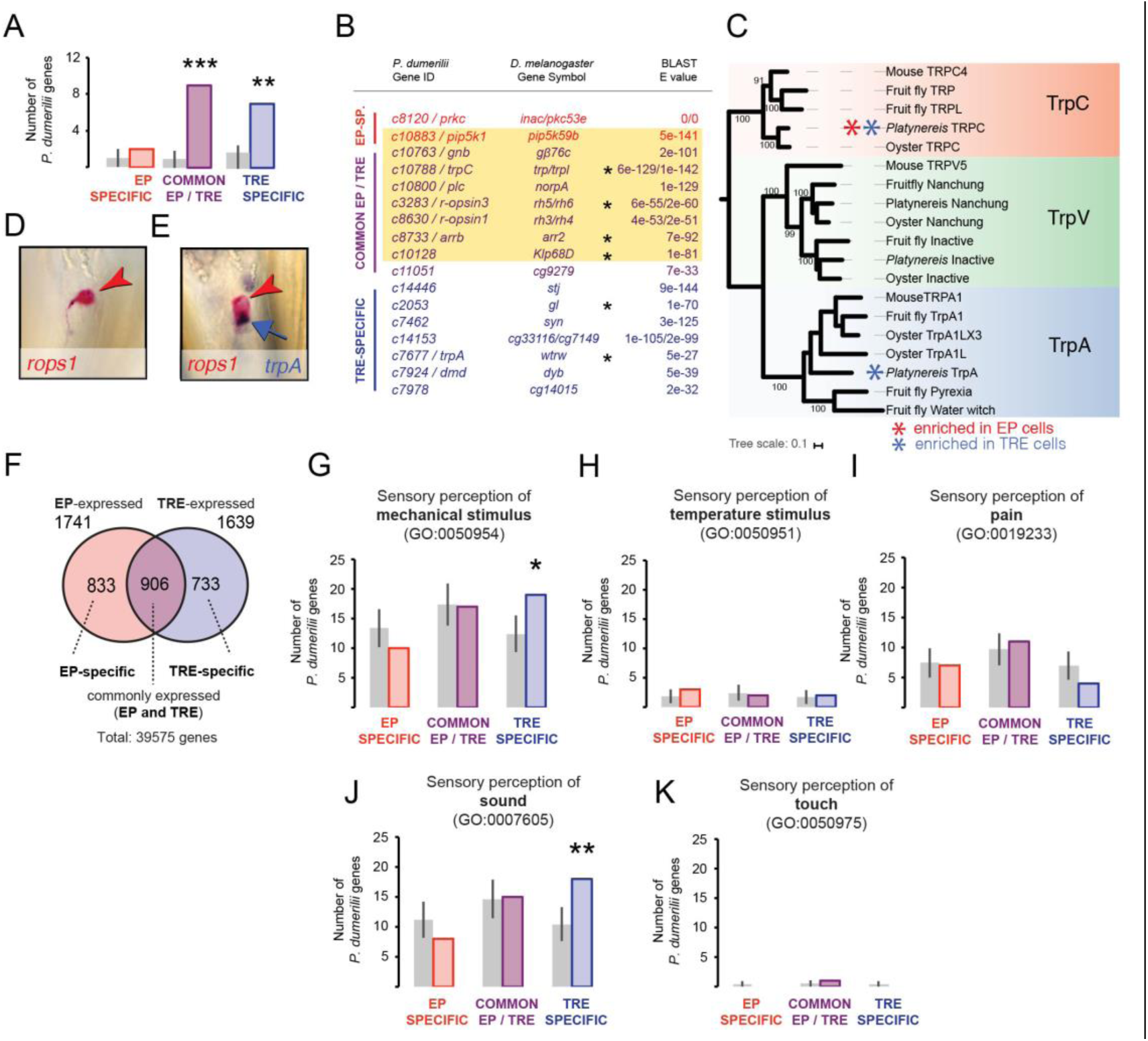
TRE cells also share a mechanosensory signature. **(A,B)** Comparison of *P. dumerilii* EP- and TRE-enriched genes with *D. melanogaster* Johnston Organ (JO)-enriched genes. For comparison with *D. melanogaster* EP-enriched genes see **Fig.3-figure supplement 1**. **(A)** Number of EP-specific (red), common EP- and TRE-enriched (purple) or TRE-specific (blue) *P. dumerilii* genes overlapping with *D. melanogaster* JO-enriched genes. Grey bars show the average number (± standard deviation) of TRE-specific, common EP- and TRE-enriched or TRE-specific *P. dumerilii* genes overlapping with randomly-selected sets of *D. melanogaster* genes. ** p < 0.01. *** p < 10^-4^. **(B)** List of the overlapping genes indicated in (A). Each gene in the “*P. dumerilii* Gene ID” column indicates the best *P. dumerilii* Blast hit of the listed *D. melanogaster* gene. The yellow shading indicates genes that are part of the *D. melanogaster* phototransduction pathway. Asterisks indicate genes relevant for auditory JO function [15]. **(C)** Molecular phylogeny of Transient receptor potential channel (Trp) orthologs showing the assignment of the joint EP/TRE-enriched TRPC channel, and the *Platynereis* TrpA ortholog expressed in the TRE cells; **(D,E)** Specific co-expression of *Platynereis r-opsin1* (D,E, red; red arrowheads) and *trpA* (E, purple; blue arrow) in TRE cells, reflecting one of various TRE markers shared with mechanosensory cells (see **Fig.2-figure supplement 4,5**); caudal views, distal to the top. **(F)** Number of genes expressed in EP and/or TRE cells. **(G–K)** Number of EP-specific (red), common EP-/TRE- expressed (purple) or TRE-specific (blue) *P. dumerilii* genes overlapping with *Mus musculus* genes involved in sensory perception of mechanical stimulus (G), sensory perception of temperature stimulus (H), sensory perception of pain (I), sensory perception of sound (J), or sensory perception of touch (K). For list of overlapping genes indicated in (G) see **Fig.3-figure supplement 2**. For list of overlapping genes indicated in (J) see **Fig.3-figure supplement 2** (yellow shading). Grey bars show the average number (± standard deviation) of TRE-specific, common EP-/TRE-enriched or TRE-specific *P. dumerilii* genes overlapping with randomly-selected sets of *Mus musculus* genes. * p < 0.05. ** p < 0.01.

Given the well-established function of JO neurons as mechanosensory cells, we next investigated whether the additional, statistically significant overlap between JO-specific genes and TRE-specific genes reflected any shared mechanosensory signature. Among the 7 JO-specific genes overlapping with TRE-specific genes, 2 were shown [15] to be required for the normal response of JO neurons to mechanical stimuli (*gl* and *wtrw;* matching *Platynereis trpA/c7677;* **Fig. 3C-E**), adding to the 4 (of 9) specifically shared genes from the joined TRE/EP set with known mechanical functions (asterisks in **Fig. 3B**), whereas the other 5 have not been tested for mechanosensory functions. In order to compensate for this lack of functional information, we also performed a comparison with mouse, where the largest number of genes involved in mechanosensation is known. We systematically determined putative *P. dumerilii* homologs of all mouse genes assigned to the gene ontology (GO) category “sensory perception of mechanical stimulus” (GO:0050954), and then assessed their overlap with EP or TRE expressed genes (**Fig. 3F,G**). Indeed, these homologs are significantly overrepresented in the TRE-specific signature (p=0.029; **Fig. 3G**). Similar analyses with GO categories for other sensory perception modalities associated to the JO, such as “sensory perception of temperature stimulus” (GO:0050951) and “sensory perception of pain” (GO:0019233), showed no statistically significant results (**Fig. 3H,I**).

A closer analysis of those mouse mechanosensory genes whose bristleworm counterparts are expressed in TRE cells (**Fig. 3-figure supplement 2**), points at a gene signature shared between TRE cells and mouse inner ear hair (IEH) cells: 18 out of the 19 TRE-specific gene homologs have reported effects on hearing function in the mouse (yellow shading in **Fig. 3-figure supplement 2**). Notably, *whrn, dnm1*, *atp8b1*, *myo3a, chrna9* and *tecta* have been shown to be required for vertebrate IEH cell function [36–40]; *sox2* and *jag2* are known to be required for development of vertebrate IEH cells [41, 42]; *crym*, *serpinb6a* and *myh14* lead to hearing loss when mutated in mammals [43–46]. Again, we assessed the specificity of this finding by systematically comparing the overlap with mouse genes involved in distinct modalities of mechanosensation, confirming a statistically significant overlap between mouse hearing genes and *P. dumerilii* TRE-specific genes (p=0.0087; **Fig. 3J**), whereas other mechanosensory modalities yielded no statistically significant results (**Fig. 3K**). Even though additional functions, unrelated to mechanosensation, are known for some of the above genes, these statistical results strongly argue for a gene signature specifically shared between *P. dumerilii* TRE cells and mouse IEH cells.

The shared mechanosensory transcriptome signature of *P. dumerilii* TRE cells, *Drosophila* JO neurons and mouse IEH cells is consistent with the possibility that TRE cells retain a combination of mechano- and photosensory molecular features, as they were previously suggested to form a likely ancestral protosensory state [19,47,48]. A deeper molecular relationship between these cells is further reinforced by the observation that the worm’s TRE cells differentiate out of a territory that expresses the gene encoding for the transcription factor Pax2/5/8 [17]. Similarly, differentiation of JO neurons and mouse IEH cells requires respective *Drosophila* (*spa*) and mouse (*pax2*) orthologs. Similarly, TRE cells have been linked to expression of *brn3/pou4f3* [17], a *Platynereis* ortholog of the vertebrate *pou4f3* gene demarcating the neuromasts of the fish lateral line [49], a set of mechanosensory structures that also express fish r-Opsin orthologs [17].

### *P. dumerilii* r-Opsin1 mediates blue light reception through Gα_q_ signaling

As a prerequisite for a deeper analysis of the function of r-Opsin1 in the TRE cells of *P. dumerilii*, we next set out to characterize the photosensory properties of this Opsin. A distinctive feature of r-Opsin phototransduction cascade is the coupling and light-dependent activation of the Gα_q_ protein by r- Opsin [50]. Amphioxus, chicken and human Melanopsins – orthologs of *Drosophila* r-Opsins – have all been shown to elicit intracellular calcium increase in response to light, and that all of these are capable of activating the Gα_q_ protein in a light-dependent manner [51]. Given that *P. dumerilii* r-Opsin1 is an ortholog of *Drosophila* r-Opsins and chordate Melanopsins [2, 52], we tested if *P. dumerilii* r-Opsin1 can also activate Gα_q_ signaling upon light exposure by employing a cell culture second messenger assay [51] (see Methods). *P. dumerilii r-opsin1*-transfected HEK293 cells exhibited a significant response to light exposure, similar to the human Melanopsin (positive control) (**Fig. 4A**). By contrast, using corresponding assays for Gα_s_ or Gα_i/o_ activation [51, 53], we detected no activation of either Gα_s_ (**Fig.4-figure supplement 1A**) or Gα_i/o_ (**Fig.4-figure supplement 1B**) by *P. dumerilii* r-Opsin1. This indicates that *Platynereis* r-Opsin1 specifically activates Gα_q_, similar to *Drosophila* r-Opsins.

**Fig. 4.**
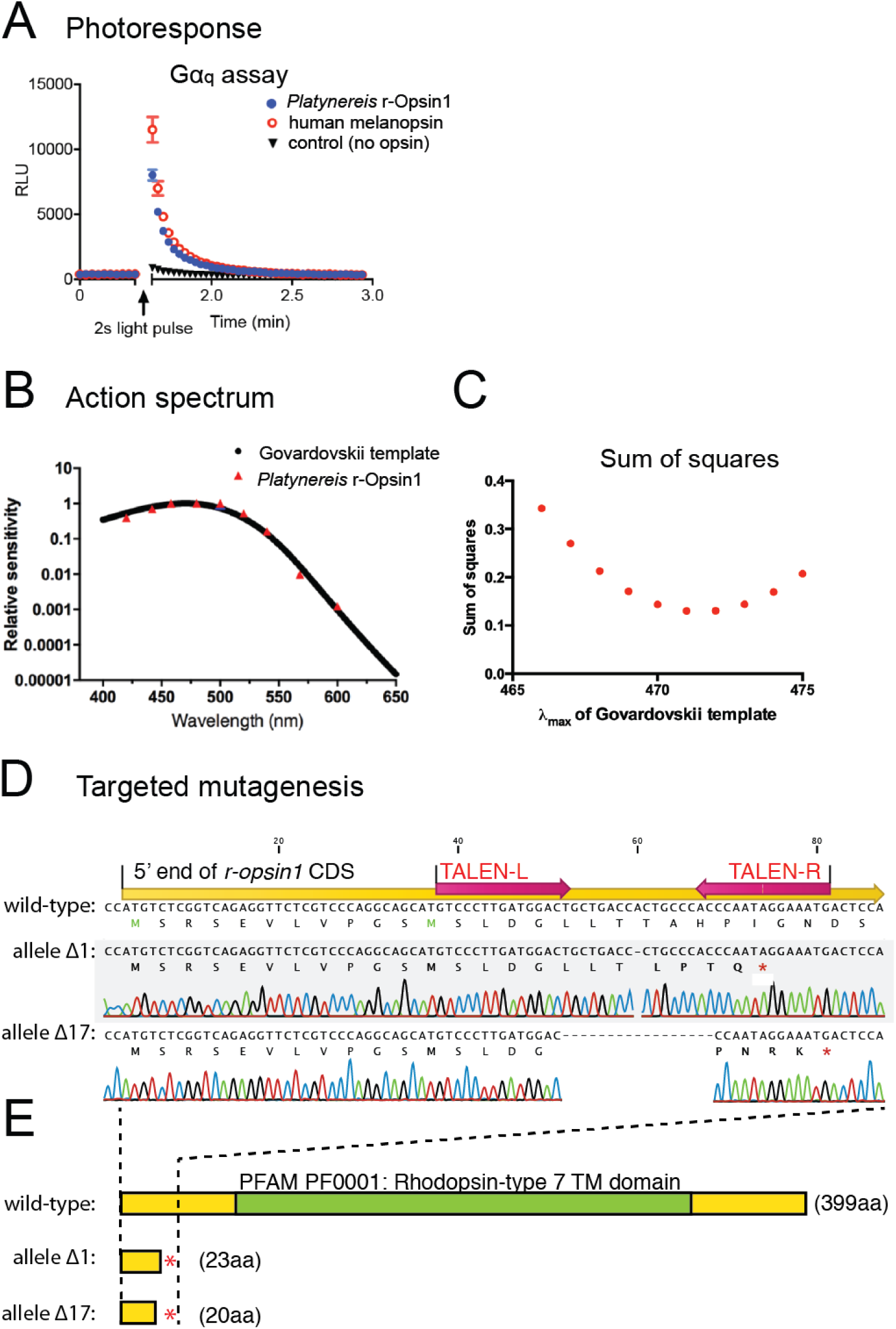
Action spectrum and targeted deletion of *Platynereis* r-Opsin1, a Gα_q_ -coupled blue-light photosensor. **(A)** Gα_q_ bioassay, showing an increase in luminescent reporter signal for calcium increase after 2s white light exposure in cells transfected with *Platynereis r-opsin1*. The increase in luminescent reporter signal is similar as when cells are transfected with the positive control human melanopsin. n= 3 independent experiments in all cases. For Gα_s_ and Gα_i/o_ assays see **Fig.4-figure supplement 1A,B**. **(B)** Action spectrum of r-Opsin1 (based on light spectra and irradiance response curves shown in **Fig.4-figure supplement 1C-F**), fit with a Govardovskii curve visual template obtained with a λ_max_ of 471 nm. **(C)** Plotted sum of squares between action spectra and Govardovskii templates at varying λ_max_, revealing a minimum for λ_max_ of 471 nm. **(D)** Targeted mutagenesis of *Platynereis r-opsin1*. Nucleotide alignment between the 5’ ends of the wild-type (top) and mutant alleles for *r-opsin1*. In the wild-type sequence, positions of the coding sequence (yellow), and of the TALE nuclease binding sites (red arrows) are indicated. Allele Δ1 contains a single nucleotide deletion, allele Δ17 lacks seventeen nucleotides; both lead to premature stop codons (marked as red asterisks in the corresponding translations). **(E)** A comparison of the encoded proteins (protein lengths indicated in brackets) reveals that alleles Δ1 and Δ17 lack the complete 7-transmembrane domain (green, PFAM domain PF0001) including the critical lysine residue for retinal binding, strongly predicting the alleles as null alleles.

The relative responsiveness of a photoreceptor cell to different wavelengths of light is a fundamental determinant of its sensory capabilities. We therefore next determined the spectral sensitivity of *P. dumerilii* r-Opsin1, using our HEK293 cells second messenger assay to measure changes in calcium concentration in response to near monochromatic stimuli spanning the visible spectrum (**Fig.4-figure supplement 1C-F**). The EC_50_ values (irradiance required to elicit 50% response; see **Fig.4-figure supplement 1F**) of sigmoidal dose response curves were converted to a relative sensitivity and fitted with an opsin:retinaldehyde pigment template function [54]. The optimal λ_max_ for the template was determined by least squares as 471 nm (**Fig. 4B,C**). *P. dumerilii* r-Opsin1 therefore maximally absorbs light in the blue range, similar to other r-Opsin orthologs, such as human Melanopsin, which exhibits a λ_max_ of around 480 nm [51].

In summary, the presence of all components of the r-Opsin phototransduction pathway in TRE cells, and our demonstration that *P. dumerilii* r-Opsin1 is capable of activating Gα_q_, strongly suggests that EP and TRE cells can respond to light.

### Mutation of *r-opsin1* affects TRE-specific, light-dependent expression of an Atp2b calcium channel involved in hearing

In order to gain insight into the function of r-Opsin1 in TRE cells, we generated two independent *r-opsin1* alleles (*r-opsin1^Δ1^* and *r-opsin^Δ17^*) in the background of the *pMos{rops::egfp}^vbci2^* strain by TALENs [20], resulting in premature stop codons in the 5’ coding region of the *r-opsin1* gene (**Fig. 4D,E**). Founders were outcrossed to wild-type worms (PIN and VIO strains). Subsequently, trans heterozygous individuals (*r-opsin1^Δ1/Δ17^*; *pMos{rops::egfp}^vbci2 +^*) were used to systematically analyze the molecular profile of EGFP-positive head and trunk cells as described above. Sampling from related EGFP-positive non-mutant specimens (*r-opsin1^+/+^*; *pMos{rops::egfp}^vbci2 +^*) served as controls to match mutant vs non-mutant profiles. Based on our spectral sensitivity results for r-Opsin1, specimens were kept under monochromatic blue light (∼470 nm, i.e. the λ_max_ of r-Opsin1), for 3 – 5 days until dissociation for FACS.

We next identified the genes differentially expressed between the EP or TRE cells of mutant vs non-mutant worms using the EdgeR algorithm. Genes with an FDR < 0.05 were considered significantly differentially expressed. We then focused on the *P. dumerilii* homologs of all mouse hearing genes that were expressed in either EP or TRE cells of mutant or non-mutant worms. In the EP cells, none of these candidate genes was significantly differentially expressed between mutant and non-mutant worms. By contrast, one gene (*atp2b/c7424*, *P. dumerilii* homolog of mouse *atp2b2*; **Fig.5-figure supplement 1A**, red arrowhead) was significantly depleted in mutant TRE cells compared to wild-type cells (FDR = 0.010; **Fig. 5A**). The specificity of this regulation is further supported by the fact that none of the identified phototransduction components were changed.

**Fig. 5.**
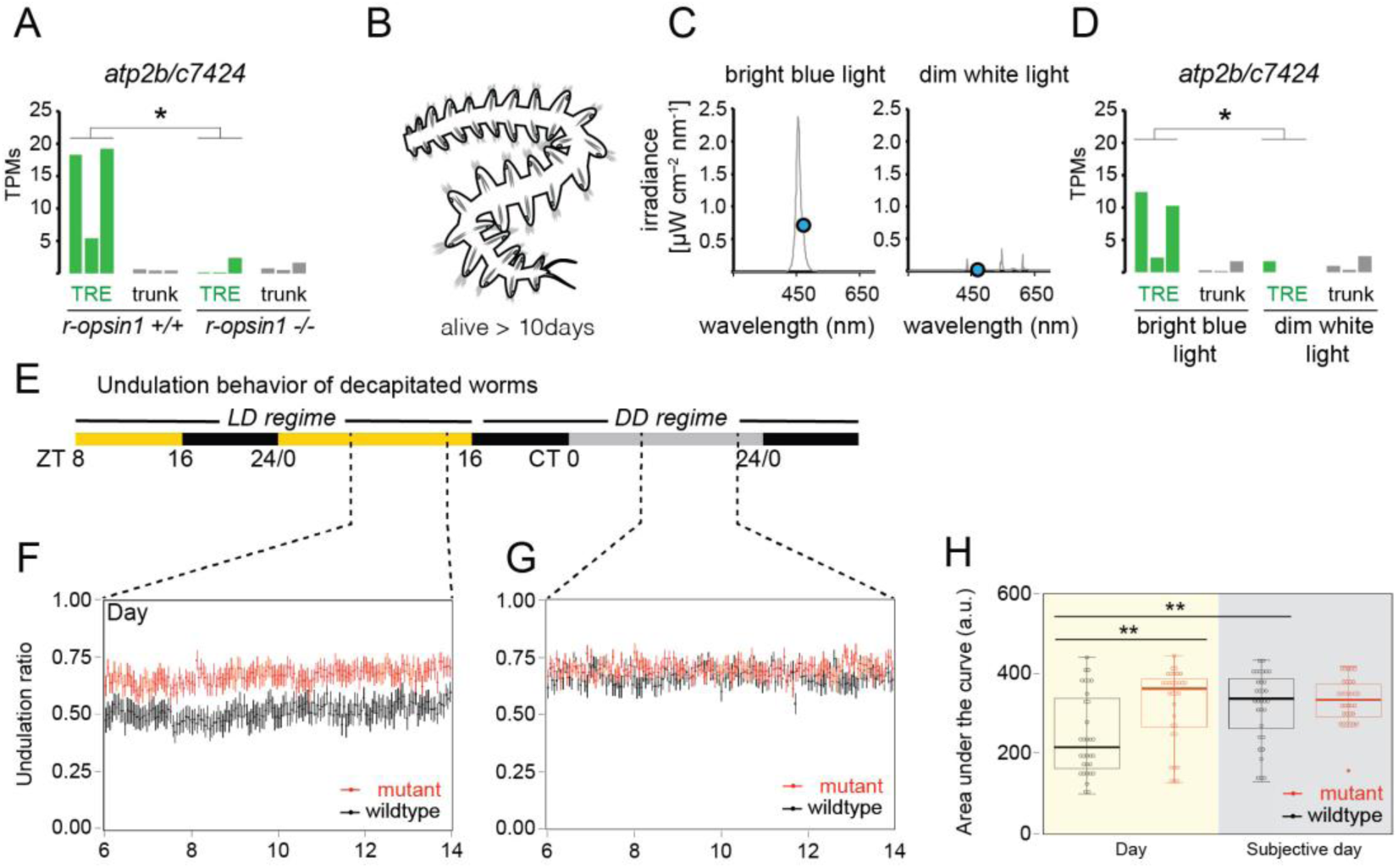
*r*-*opsin1* mediates blue-light modulation of TRE signature and undulation behavior. **(A)** *atp2b/c7424* expression levels (in TPMs) in individual replicates of *r-opsin1+/+* and *r-opsin1-/-* worms cultured for 3 -5 days in bright blue light. For Atp2b2 phylogeny see **Fig.5-figure supplement 1**. **(B)** Scheme of decapitated worm trunks as used in experiments C-H, that survive for up to 14days. **(C)** Spectral profile of bright blue and dim white light. The blue dot indicates the irradiance at 471 nm (λ_max_ of *P. dumerilii* r-Opsin1). **(D)** *atp2b/c7424* expression levels (in TPMs) in individual replicates of decapitated worms cultured for 3 -5 days in bright blue light or dim white light. **(E-H)** Undulation behavior of decapitated worms. **(E)** Light regime. Black portions of the horizontal bar indicate “night” (light off), yellow portions indicate “day” (light on), and grey portions indicate “subjective day” (light off during “day” period). ZT: Zeitgeber Time. CT: Circadian Time. **(F, G)** Undulation ratio during “day” (F) and “subjective day” (G). Each black (red) point represents the mean of all wild-type (mutant) worms within a 3-minute window, and vertical bars represent the standard error of the mean (n=32 for each genotype, distributed among three independent experiments). For reliability tests of the algorithm used to detect undulation behavior see **Fig.5-figure supplement 3**. **(H)** Area under the curve obtained from the undulation ratios shown in (F)(yellow background; “Day”) and (G)(grey background; “Subjective day”). Circles indicate data corresponding to individual worms. Box plots indicate the median (thick horizontal line), the 50% quantile (box) and 100% quantile (error bars). Filled circle indicates an outlier (as determined by the boxplot function of the ggplot R package). * p-value < 0.05; ** p-value < 0.01 (Wilcoxon rank sum and signed rank tests). For behavioral responses to strong light of *r-opsin+/+* and *r-opsin-/-* trunks see **Fig.5-figure supplement 2**.

The *atp2b2* gene encodes a plasma membrane calcium-transporting ATPase, which is expressed in the stereocilia of mechanosensory cells of the murine cochlea and vestibular system [55]. Homozygous *atp2b2* mutant mice show balance deficits and are deaf, while heterozygous mutants show partial loss of auditory ability [56]. These differential effects caused by different genetic dosages of *atp2b2* are consistent with the possibility that regulation of *atp2b2* expression could be a natural mechanism to modulate mechanosensory cell function. In support of an ancestral role of the plasma membrane calcium-transporting ATPase gene family in modulating neuronal sensitivity, the single *Drosophila* representative of this family, *pmca* (**Fig.5-figure supplement 1A**, light blue arrowhead), modulates the thermal sensitivity of motor neurons [57]. Its *C. elegans* ortholog, *mca-3* (**Fig.5-figure supplement 1A**, green arrowhead), modulates touch sensitivity of the touch neurons [58]. Furthermore, in zebrafish, *atp2b2* (**Fig.5-figure supplement 1A**, dark blue arrowhead) is highly enriched in the *r-opsin*-expressing mechanosensory neuromasts of the lateral line (**Fig.5-figure supplement 1B**), consistent with a potential mechanosensory function of the gene in this organism.

Since the expression levels of *atp2b/c7424* in *Platynereis* TREs depend on *r-opsin1* function, we wondered if they might also depend on illumination. Decapitated worm trunks are functionally relatively autonomous, maintain their ability to crawl and swim, aspects of their rhythmicity and live for up to two weeks (**Fig. 5B**) [59]. We cultured decapitated trunks of pMos{rops::egfp}^vbci2^ worms for 3-5 days in two distinct light conditions: (i) bright monochromatic blue light (∼470 nm); and (ii) very dim white light (**Fig. 5C**). Light intensity at the λ_max_ of r-Opsin1 (471 nm) was ∼40 fold reduced in the dim light condition compared to bright blue light (blue circles in **Fig. 5C**). TRE cells were isolated and profiled as before. Statistical analysis showed that *atp2b/c7424* is significantly downregulated in dim light conditions as compared to bright light (p=0,02; Wilcoxon rank sum test; **Fig. 5D**), similar to the downregulation observed in mutant worms (p=0.02; Wilcoxon rank sum test; **Fig. 5A**). These results indicate that blue light levels modulate *atp2b/c7424* expression levels in TRE cells, and suggest that the light-dependent modulation is majorly mediated by r-Opsin1.

### r-Opsin1 mediates a light-dependent modulation of undulation frequency

Given the functional relevance of *atp2b2* gene dosage in mammalian hearing, and its enrichment in zebrafish mechanosensory cells known to express *r-opsin* orthologs *opn4xb* and *opn4.1* [17] (**Fig.5-figure supplement 1B**), we hypothesized that the regulation of *atp2b/c7424* in TRE cells might correlate with altered mechanosensory abilities. We therefore set out to test the impact of changed light conditions as well as different genotypes on worm behavior. Classical studies have provided evidence for the existence of several classes of mechanosensory cells in parapodia of annelids. These include stretch-sensitive flap receptors, bristle receptors and acicular receptors [60, 61]. A plausible function of these receptors is to fine-tune motor patterns associated with directional (crawling) or stationary (undulation) movements that require coordinated activity by individual segments.

In a first experiment to assess the possible requirement of *r-opsin1* for coordinated segmental movements, we assessed the crawling movement exhibited by decapitated trunks when stimulated by a focal bright light stimulus [17]. Transheterozygous *r-opsin1^Δ1/Δ17^* individuals clearly responded to such stimuli, but exhibited a significantly reduced net distance when compared to wild-type animals (p=0.02; Wilcoxon rank sum test; **Fig.5-figure supplement 2**). Whereas this result is consistent with the notion that r-Opsin1 is involved in the correct execution of motor movements, the experiment does not discriminate between r-Opsin1 triggering the response and/or modulating its motor execution.

We therefore decided to investigate a very regularly performed behavior that does not require light as a stimulus. Annelids from the *Platynereis* genus exhibit a stereotypical undulatory behavior that is thought to increase water flow and oxygenation [62]. The presence of this behavior in *Platynereis dumerilii* is seemingly independent of time [26], and requires a tight coordination between segments. Thus, we reasoned that if r-Opsin1 in the segmentally arranged TRE cells plays a role in the modulation of motor movements, this behavior presents a good test. We recorded the movement of *r-opsin1* mutant and wild-type trunks of de-capitated worms for five consecutive days, using a previously established infrared video system [63]. Concerning visible light conditions, during the first 1.5 days, recorded worms were kept under a light/dark (LD) regime of 16:8 hours, followed by constant darkness (DD, **Fig. 5E**). We then established a deep-learning-based quantitative behavioral approach to analyze the resulting movies. We trained a neural network to detect 7 different body positions: *jaws, body1-body5*, *tail* (**Fig.5-figure supplement 3A**) across the total length of each movie. Next, we analyzed 10-second intervals of the movie to identify oscillatory behavior of the *body1* through *body5* points, using a periodogram algorithm, categorizing each interval into undulatory or non-undulatory behavior. This automated analytical setup was benchmarked against human observations of a portion of the movies (**Fig.5-figure supplement 3B,C**). It allowed us to systematically determine the ratio of time that specimens spent undulating compared to the overall time (**Fig. 5F,G**). In turn, this permitted us to compare both the effect of *r-opsin1* mutation (red graphs in **Fig. 5F,G**) to wild-types (black graphs in **Fig. 5F,G**) and the effect of illumination (day, **Fig. 5F**) compared to darkness (subjective day, **Fig. 5G**) in equivalent windows of circadian time.

Analyses on a total of 64 trunks revealed that wild-type (black graphs) exhibited a light-dependent modulation of the undulatory movements, which were higher during darkness (**Fig. 5F-H**). This modulation was abolished in *r-ops1-/-* worms, whose trunks exhibited equally high undulatory movements during light and dark (**Fig.5F-H**, red graphs). (Please note that in complete animals the difference between wild-type and mutants is also present, but the effect of light modulation on wild-type movements is inversed, data not shown.)

These result couple r-Opsin1 in TREs with their mechanosensory molecular signature with the light-dependent modulation of regular behavioral movements.

## DISCUSSION

Shared and distinct molecular signatures, like those we derived from FAC-sorted EP and TRE cells, are valuable substrates for inferring cell type divergence and evolution [64–66]. Our finding that both *Platynereis* TRE cells and *Drosophila* JO neurons retain a close-to-complete r-Opsin phototransduction machinery suggests as most parsimonious explanation that *r-opsin-*expressing photo- and mechanoreceptors are cell types that arose from a common ancestral cell type early in animal evolution. This notion is supported by the fact that photoreceptive and mechanoreceptive cells rely both on shared specification factors such as *atonal/Atonal2/5* and *pou4f3* [19,48,67], and by distinct ones that have arisen by gene duplication, such the Pax genes *pax6/ey* – primarily associated with eye photoreceptors – and *pax2/5/8/spa* – primarily associated with mechanoreceptive cells [19,47,67]. In accordance with this notion, development of the worm’s TRE cells, found here to exhibit a mechanosensory signature, has been linked to *brn3/pou4f3* and *pax2/5/8* [17], and cnidarian PaxB, a transcription factor that combines features of both Pax6/Ey and Pax2/5/8/Spa has been shown to be involved in the formation of the rhopalia in the Cubozoan jellyfish *Tripedalia crystophora* [67]. The rhopalia are sensory structures that combine photo- and mechanosensory functions. Our observation that EP and TRE cells also express deep homologs of the Transient receptor potential (Trp) channel family (TrpC and TrpA, respectively, **Fig. 3C**) not only adds to this concept on the level of effector molecules, but also argues that the ancient cell type already possessed one or more sensors for membrane stretch. Similarly, the close molecular relationships between mechanosensory cells of the lateral line and ear and photosensory cell types present during vertebrate development (reviewed in [68]), and the uncovered genetic links between ear- and eye defects revealed in human conditions such as the Usher syndrome [69] might also reflect such a deep homology in the specification of sensory cells.

Based on the observation that rhabdomeric Opsins appear to serve light-independent structural roles in the fly’s mechanosensory cells of the JO and ChO, it has been suggested that such light-independent, cell-mechanical roles are the ancestral function of animal r-Opsins [16, 70]. However, r-Opsins only constitute one of nine Opsin families that already existed at the dawn of bilaterian evolution, and light sensitivity is a common feature of its extant members [3]. Thus, the evolutionary hypothesis of an ancestral primary non-light sensory function of one bilaterian subgroup either implies that light sensitivity evolved independently in distinct Opsin groups, or that r-Opsins would have undergone a loss of light sensitivity prior to evolving this feature again. A more plausible explanation is that light sensitivity is an ancient feature of r-Opsins, and that the close association of r-Opsins and certain mechanosensors reflects an ancestral role of light in such cells.

Indeed, our data are consistent with a concept in which Opsins endow mechanoreceptors with the ability to tune their responses in response to environmental light conditions, on at least two levels: A first level are light- and *r-opsin1-*dependent changes in transcript levels of *atp2b/c7424*. As ATP2B2 is an ion transport ATPase, which removes Ca^2+^ from the cytoplasm, different expression levels of this enzyme can impact on the time after which a neuron will return to its resting state. Thereby, changing *atp2b* levels likely modulates signal transduction and/or refractory period of cells, resulting in overall changes in receptor sensitivity. This model is consistent with both the relevance of *r-opsin1* for tuning the undulatory behavior of trunks to ambient light conditions in the bristleworm, and the differential effects of different genetic dosages of *atp2b2* (homozygous vs. heterozygous state) in mice [56]. While we have not directly assessed the speed by which *atp2b/c7424* transcript-levels are modulated, such changes would be expected to take place on the scale of minutes to hours, thus providing a slow adjustment of signaling potential.

A second mechanism by which r-Opsins could modulate mechanosensation more acutely is provided by the photomechanical response that was uncovered by the study of *Drosophila* EP function [71]. Specifically, this model proposes that Opsin-induced, phospholipase C-mediated PIP_2_ cleavage results in a fast-propagating change in photoreceptor bilayer curvature that then triggers stretch-sensitive TRP-C channels. Thereby, photon absorption (light reception) is effectively translated into a local stretch signal as it is at the core of various mechanosensory cell types. Given that this mechanism seems to account for a canonical photoreceptive function of r-Opsin in EP cells, the conservation of *r-opsin* expression along with the respective signaling machinery suggests that Opsin activation in mechanoreceptive cells may well acutely tune the membrane curvature and thus the ability of stretch receptors to be activated.

From an ecological perspective, a light-modulatory function could effectively serve to adjust mechanosensory functions in species exposed to varying light conditions, allowing them to tune mechanoreceptive responses to ambient light. Whereas our functional results are restricted to the bristleworm model, we reason that a modulatory function as proposed here might plausibly also reflect the functionality of an ancestral “protosensory” cell [47], that could subsequently have been subfunctionalized into dedicated light sensory and mechanoreceptive cell types. From this perspective, the absence of apparent light sensitivity in *Drosophila* JO or ChO neurons likely represent secondary evolutionary processes rather than ancestral conditions. Likewise, similar principles might apply to the apparent light-independent functions of r-Opsins in chemosensory cells suggested by recent experiments in the fruitfly [72], as chemosensory cells were also noted to share molecular signature with r-Opsin light sensors before [19]. Furthermore, we note that in specific neurons of the cnidarian *Hydra magnipapillata*, the signaling pathway downstream of a distinct Opsin class (Cnidops) has been suggested to modulate the discharge of neighboring cnidocytes, a complex cell type also exhibiting sensory functions [73, 74]. It remains unclear if this link reflects parallel evolution or, alternatively, an even deeper link between Opsins and sensory cells. In either setting, however, this finding strengthens the notion that light modulation of animal mechanosensation is a fundamental principle.

Finally, our study also advances technology establishment for a “non-conventional model system” at multiples levels. First, the FACS-based protocol for cell type profiling employed here will be useful in the context of other non-conventional marine model organisms. Second, we anticipate that the automatic analyses of behavioral types by deep-learning based software tools will provide new opportunities to identify and quantify behavioral paradigms under different environmental and genetic conditions.

## ACKNOWLEDGEMENTS

We thank the members of the Tessmar-Raible and Raible groups for discussions, and Andrij Belokurov, Margaryta Borysova and Netsanet Getachew for help with worm care and genotyping at the Max Perutz Labs aquatic facility.

The research leading to these results has received funding from the European Research Council under the European Community‘s Seventh Framework Programme (FP7/2007–2013)/ERC Grant Agreement 260304 (F.R.) and ERC Grant Agreement 337011 (K.T.-R.); the research platforms ‘Rhythms of Life’ (K.T.-R., F.R., A.v.H.) and “Single-cell genomics of stem cells” (F.R.) of the University of Vienna; the Austrian Science Fund (FWF) START award, project Y413 (K.T.-R.); Austrian Science Fund (FWF) projects P28970 (K.T.-R.) and I2972 (F.R.). A.v.H and M.S. acknowledge financial support from the University of Vienna and the Medical University of Vienna. R.R. was supported by the Vienna International PostDoctoral Program for Molecular Life Sciences (VIPS).

None of the funding bodies was involved in the design of the study, the collection, analysis, and interpretation of data or in writing the manuscript.

## Figure Supplements

**Fig. 1- figure supplement 1.**
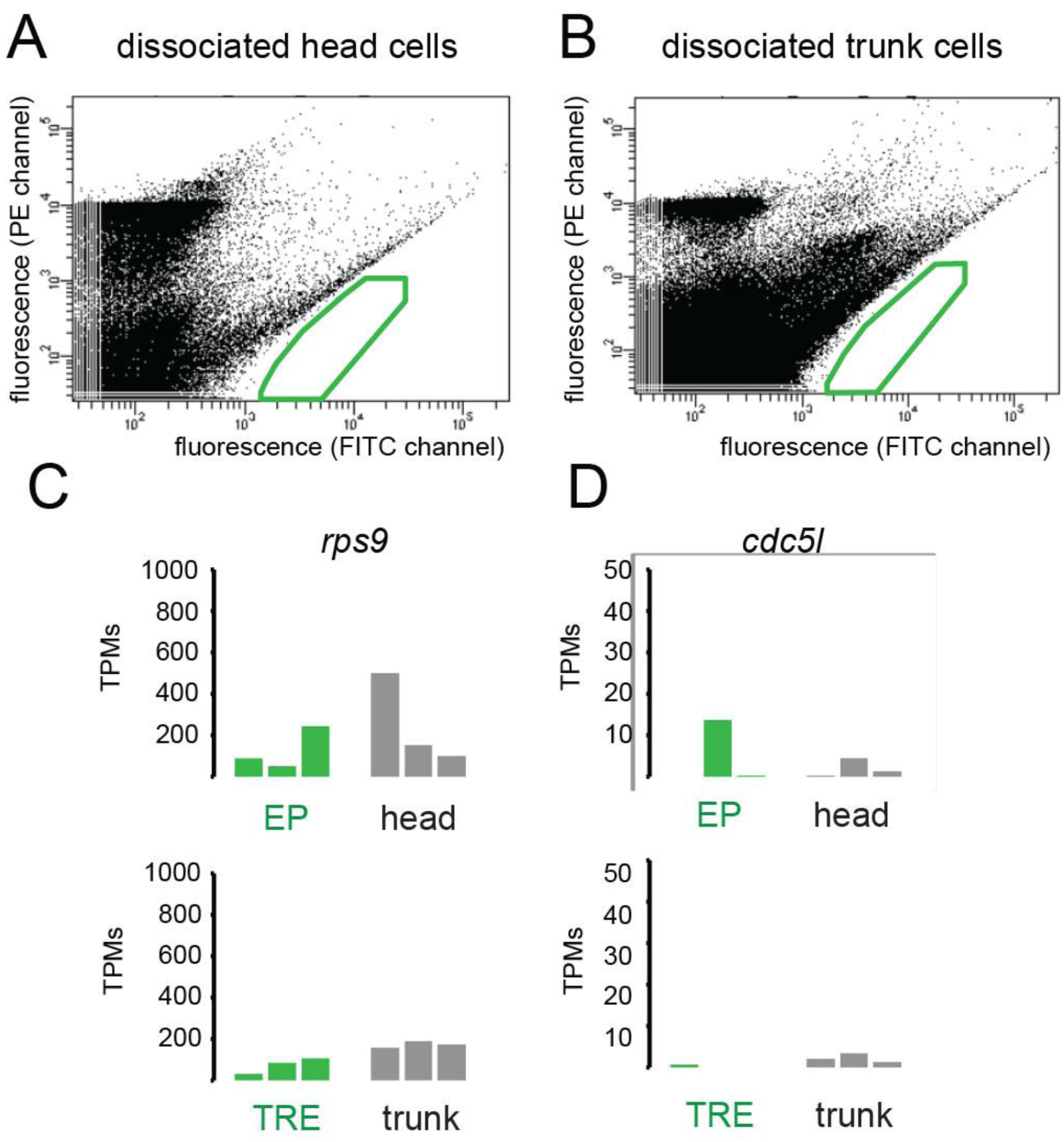
FACS profiles of dissociated cells from wild-type heads and trunks. **(A)** profile for wild-type head cells; **(B)** Profile for wild-type trunk cells. The areas boxed in green indicate gates chosen for the isolation of EGFP cells from transgenic pMos{rops::egfp}^vbci2^ individuals. **(C,D)** Lack of enrichment of reference genes *rps9* and *cdc5-like* in head- and trunk-derived libraries. Comparison of transcripts per million reads (TPMs) for the genes *rps9* (A) and *cdc5-like/cdc5l* (B) in individual replicates of EP, head, TRE, and trunk libraries. (cf. **Fig.1**).

**Fig. 2- figure supplement 1.**
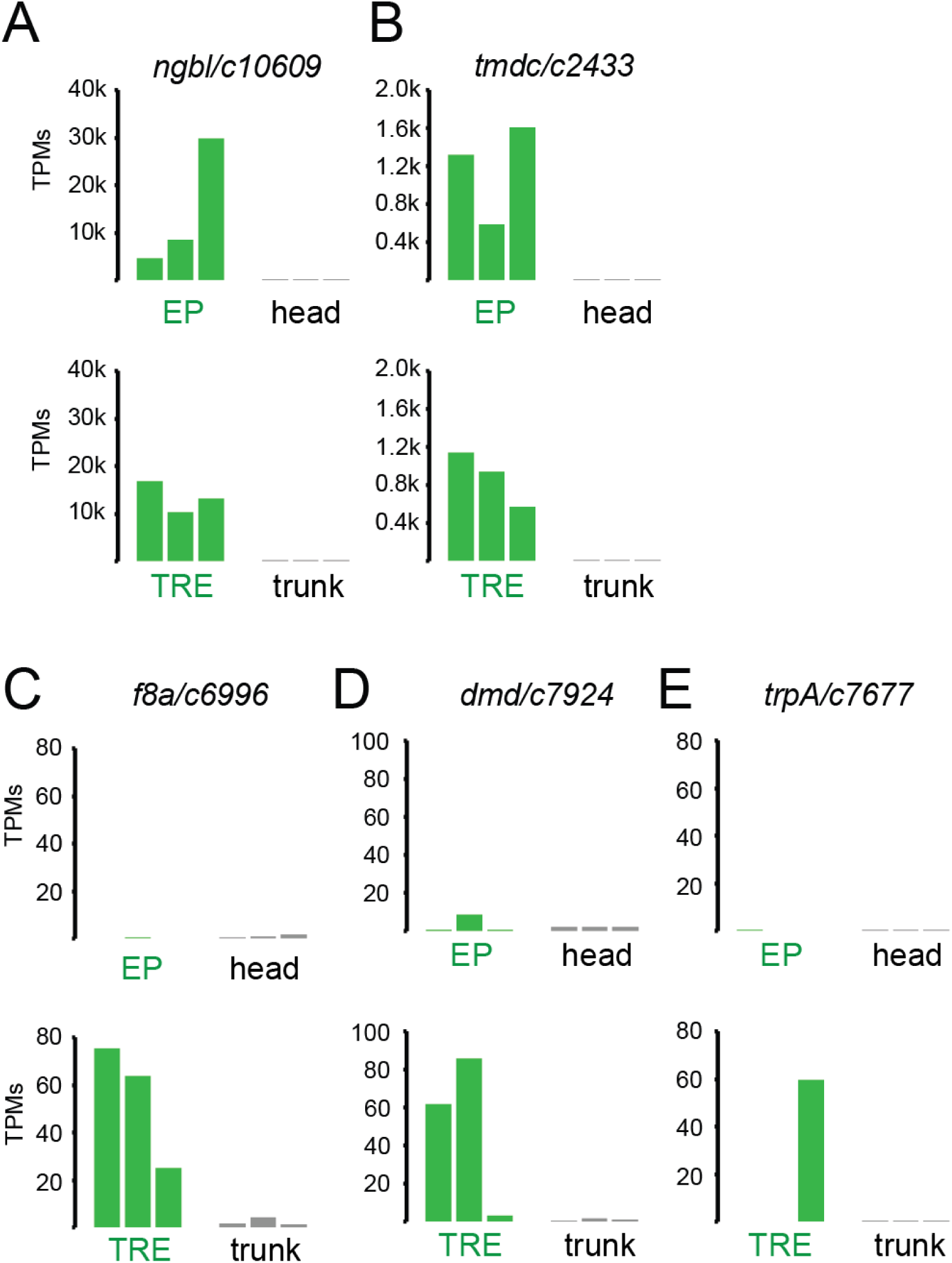
Expression levels (in TPMs, in individual replicates) of enriched genes chosen for validation. **(A,B)** Chosen genes enriched in both EP and TRE cells: *ngb/c10609* (A) and *tm/c2433* (B). **(C,D,E)** Chosen genes specifically enriched in the TRE cells: *f8a/c6996* (C), *dmd/c7924* (D) and *trpa/c7677* (E).

**Fig. 2- figure supplement 2.**
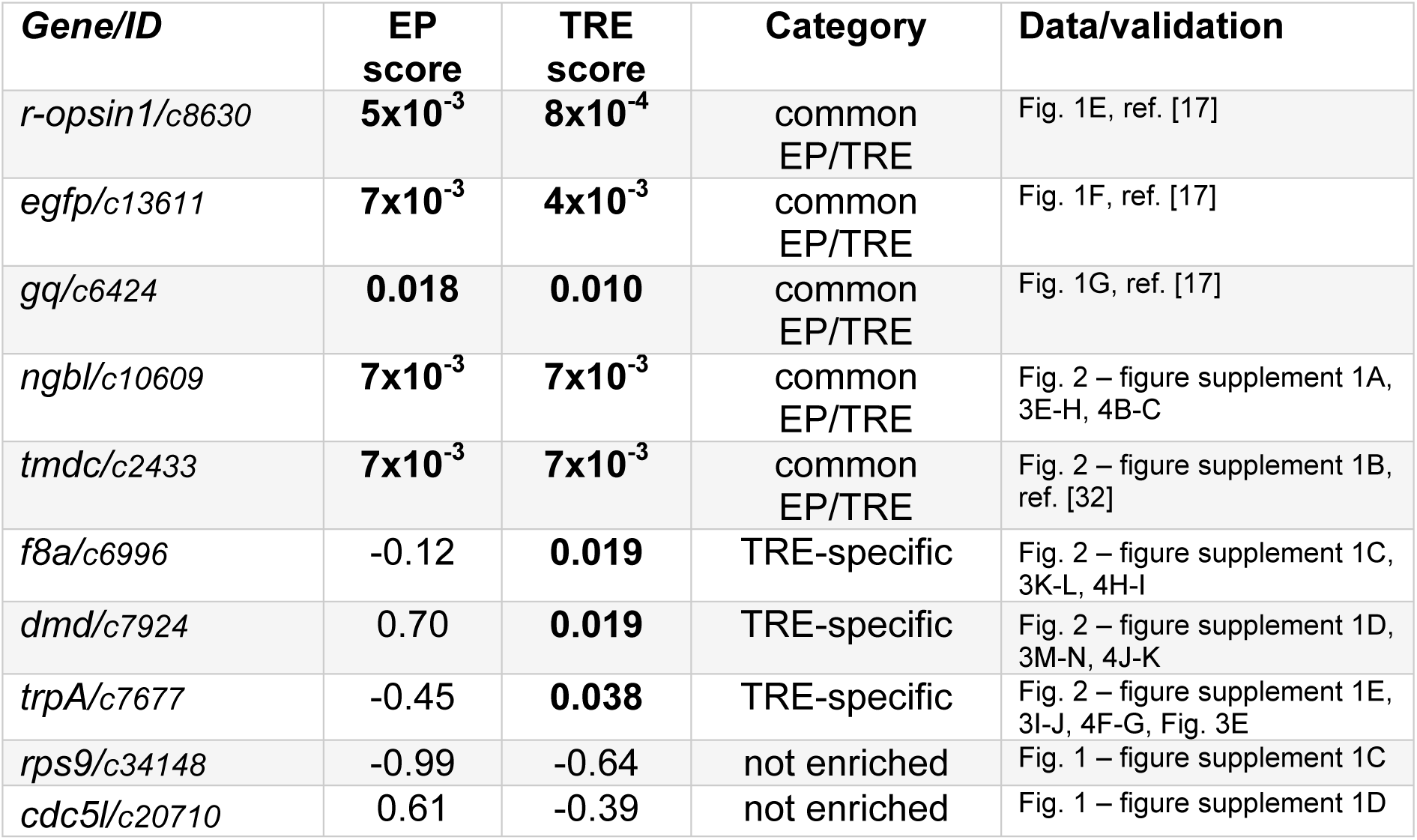
Synopsis of validated genes identified in the transcriptome profiling.

**Fig. 2- figure supplement 3.**
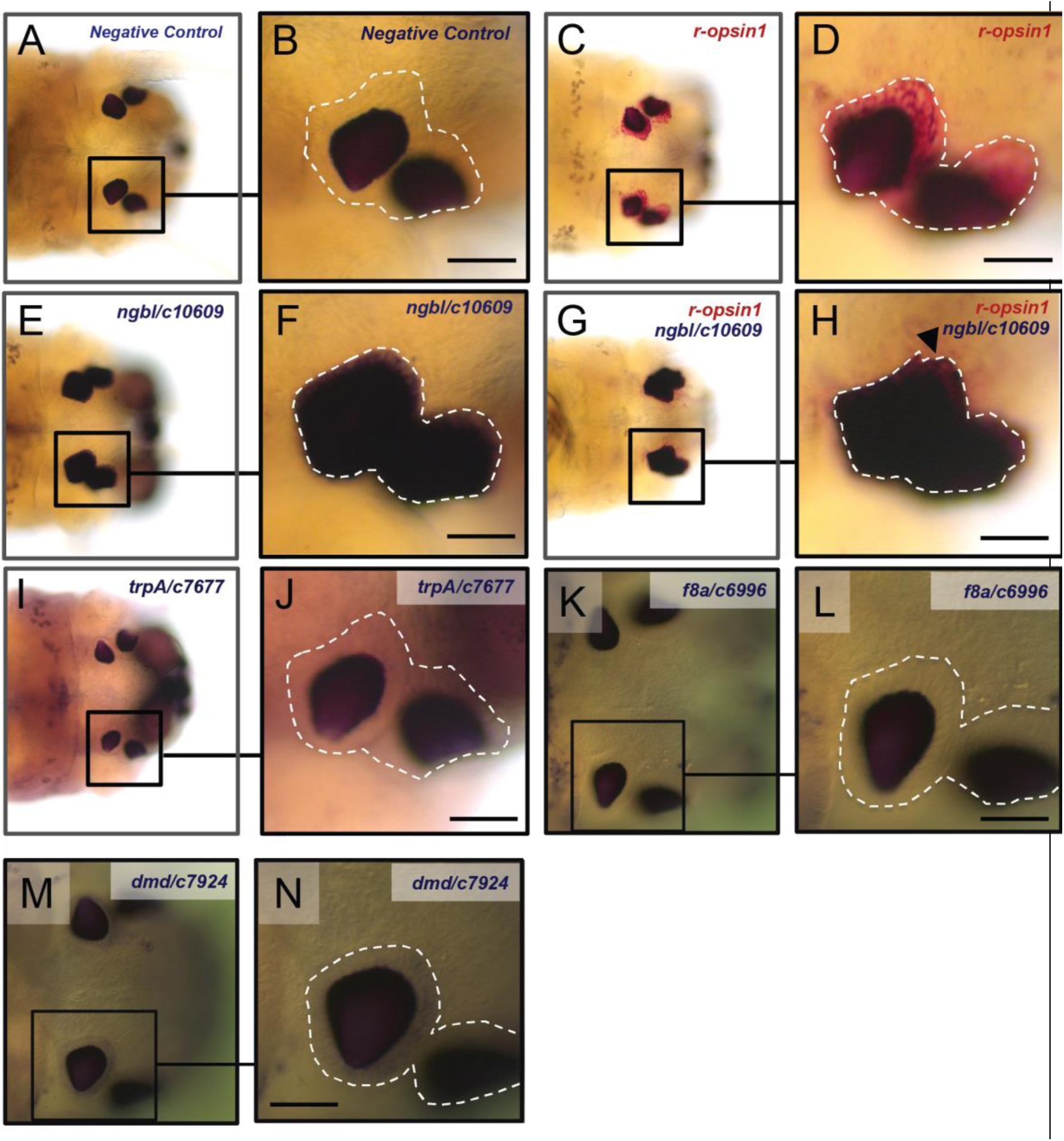
Validation of selected genes from the differential enrichment analysis (head). Low-magnification images (A,C,E,G,I,K,M) and high-magnification images (B,D,F,H,J,L,N) of single-color (A-F, I-N) or double-color (G,H) whole-mount in situ hybridization experiments using the following probes: A sense probe (A,B) showing no staining in the eyes; a probe against *r-ospin1* (C,D,G,H, red staining) showing expression in the eye photoreceptors; a probe against *ngbl/c10609* (E,F,G,H, blue staining) showing expression in the eyes that overlaps with the expression of *r-opsin1* (arrowhead in H); probes against trpA/c7677 (I,J, blue staining), *f8a/c6996* (K,L, blue staining) and *dmd/c7924* (M,N, blue staining) showing no detectable expression in the eyes. The white broken line in B,D,F,H,J,L,N demarcates the region of the EP cells (see Supplementary Text). All scale bars: 50µm.

**Fig. 2- figure supplement 4.**
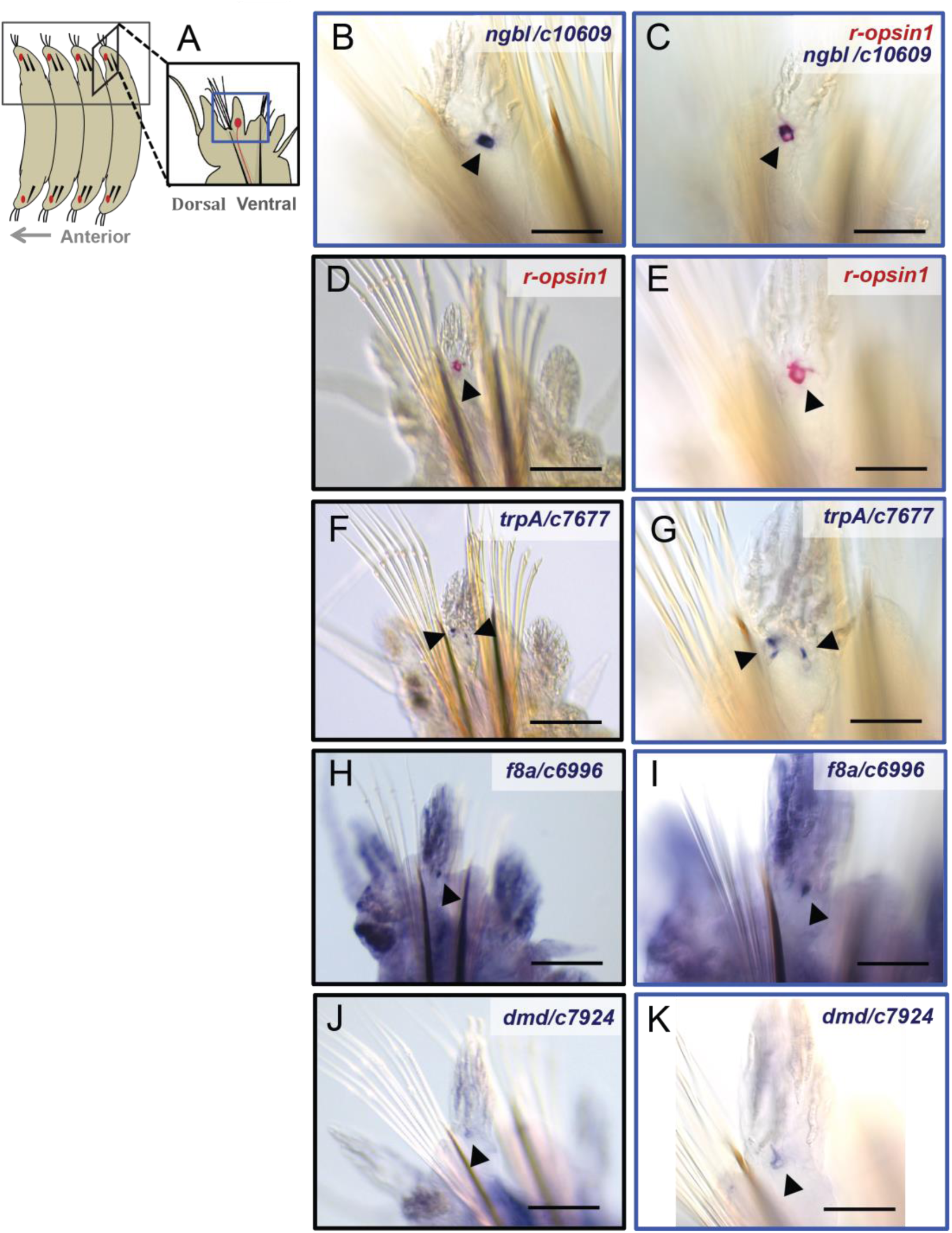
Validation of selected genes of the differential enrichment analysis (trunk). **(A)** Scheme showing the position of the TRE cells in the trunk. **(B–K)** Low magnification images (D,F,H,J) and high magnification images (B,C,E,G,I,K) of trunk WMISH (D-K) and double-WMISH (B,C) using the following probes: a probe against *r-opsin1* (C,D,E, red staining), showing expression in the TRE cell (arrowheads); a probe against *ngbl/c10609* (B,C, blue staining), showing expression in the TRE cell overlapping with the expression of *r-opsin1* (C); a probe against *trpA/c7677* (F,G, blue staining), showing expression in several spots (arrowheads) in the location of the TRE (Fig. 3D,E); probes against *f8a/c6996* (H,I, blue staining) and *dmd/c7924* (J,K, blue staining) showing expression in a single spot in a location consistent with the TRE cell. Scale bars in D,E,H,J: 100µm. Scale bars in B,C,E,G,I,K: 40µm.

**Fig. 2- figure supplement 5.**
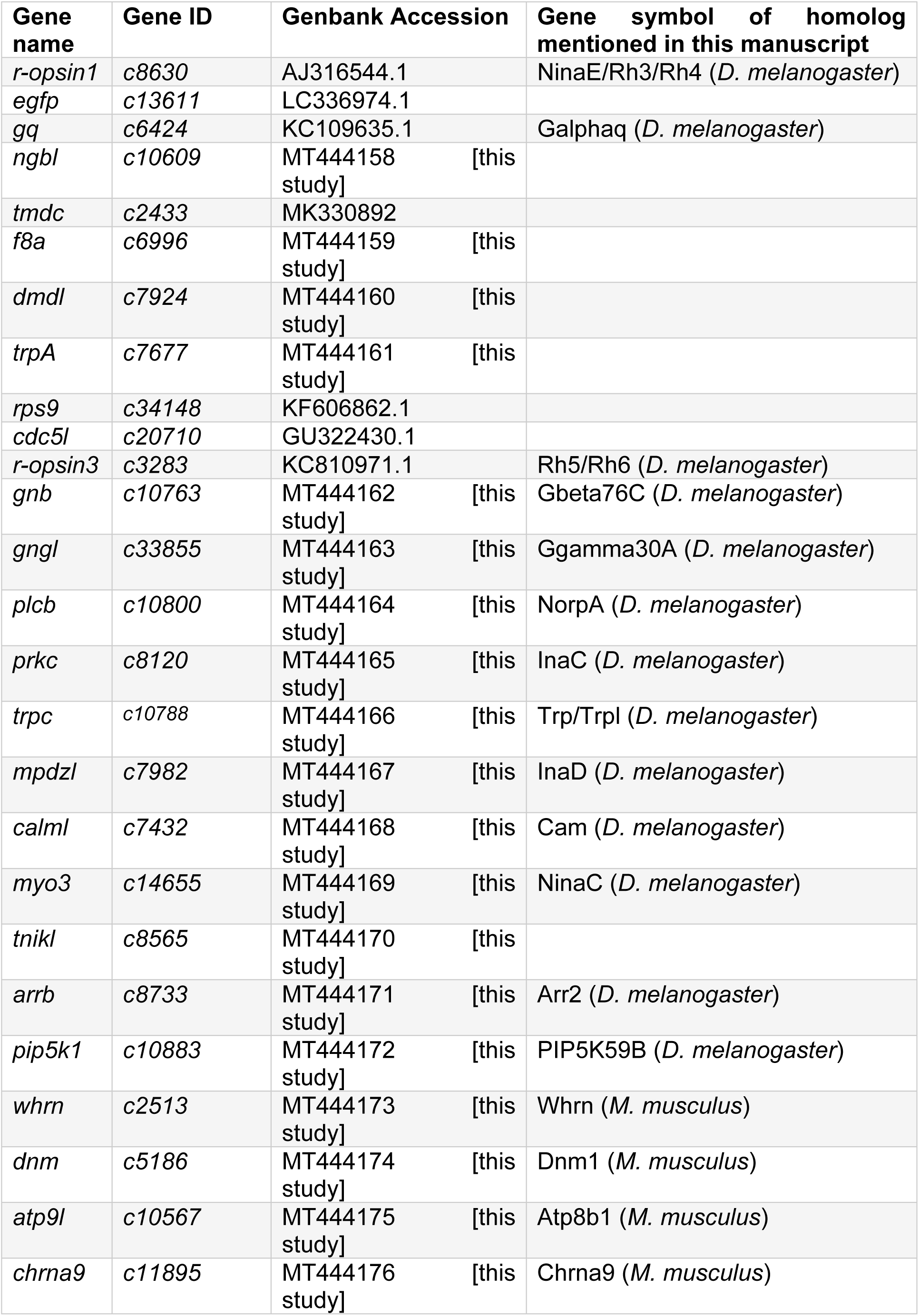
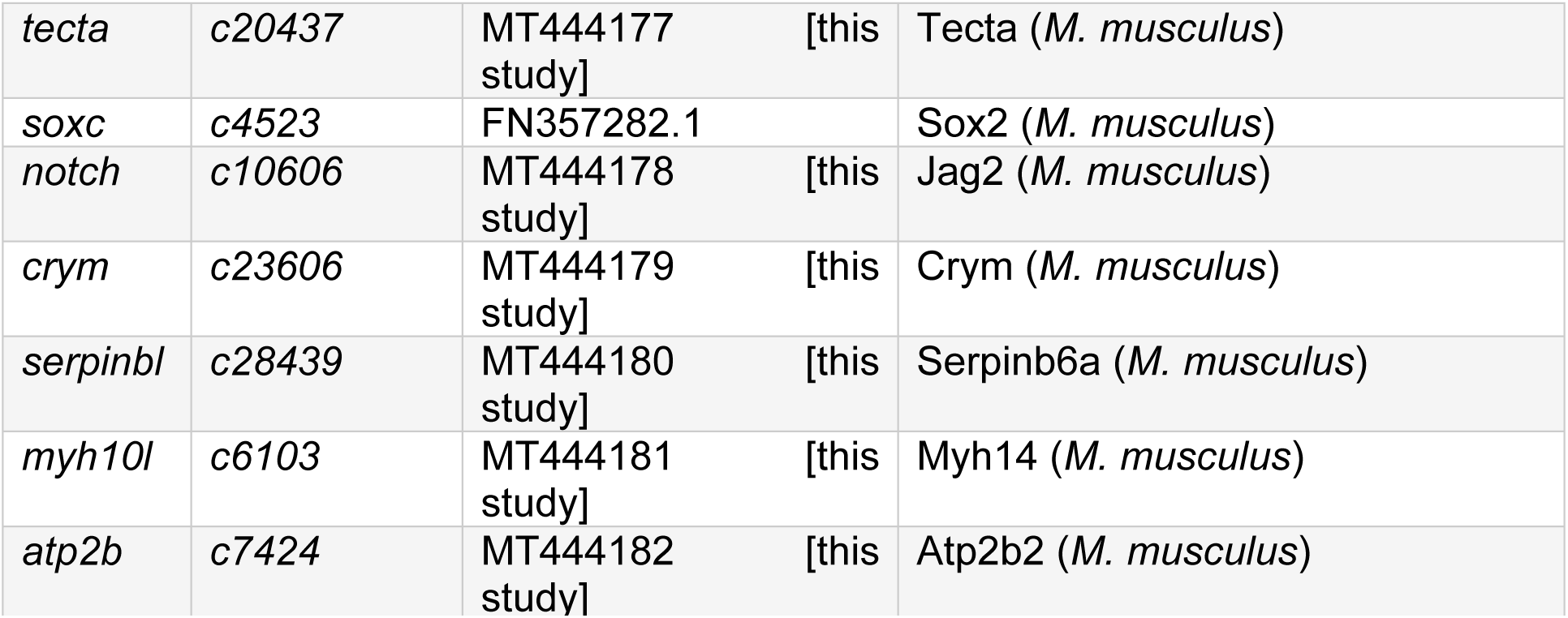
Sequence identifiers of *Platynereis* genes analyzed in this study.

**Fig. 3- figure supplement 1.**
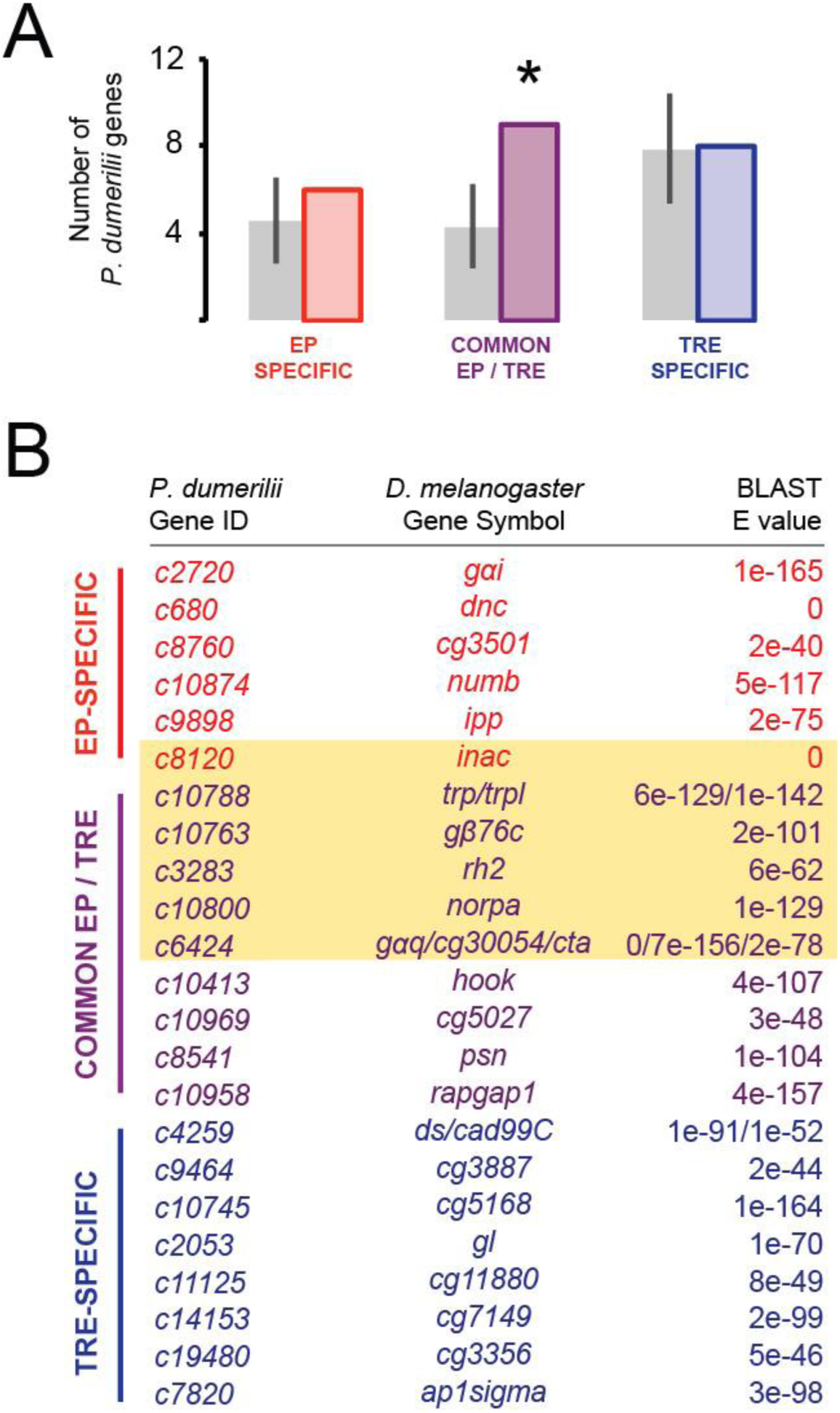
Comparison of *P. dumerilii* EP- and TRE-enriched genes with *D. melanogaster* EP-enriched genes. **(A)** Number of EP-specific (red), common EP- and TRE-enriched (purple) or TRE-specific (blue) *P. dumerilii* genes overlapping with *D. melanogaster* EP-enriched genes. Grey bars show the average number (± standard deviation) of TRE-specific, common EP- and TRE- enriched or TRE-specific *P. dumerilii* genes overlapping with randomly-selected sets of *D. melanogaster* genes. * p < 0.05. **(B)** List of the overlapping genes indicated in (A). Best *P. dumerilii* Blast hits of the corresponding *D. melanogaster* genes (middle column), as described in detail in Methods. The yellow shading indicates genes that are part of the *D. melanogaster* phototransduction pathway.

**Fig. 3- figure supplement 2.**
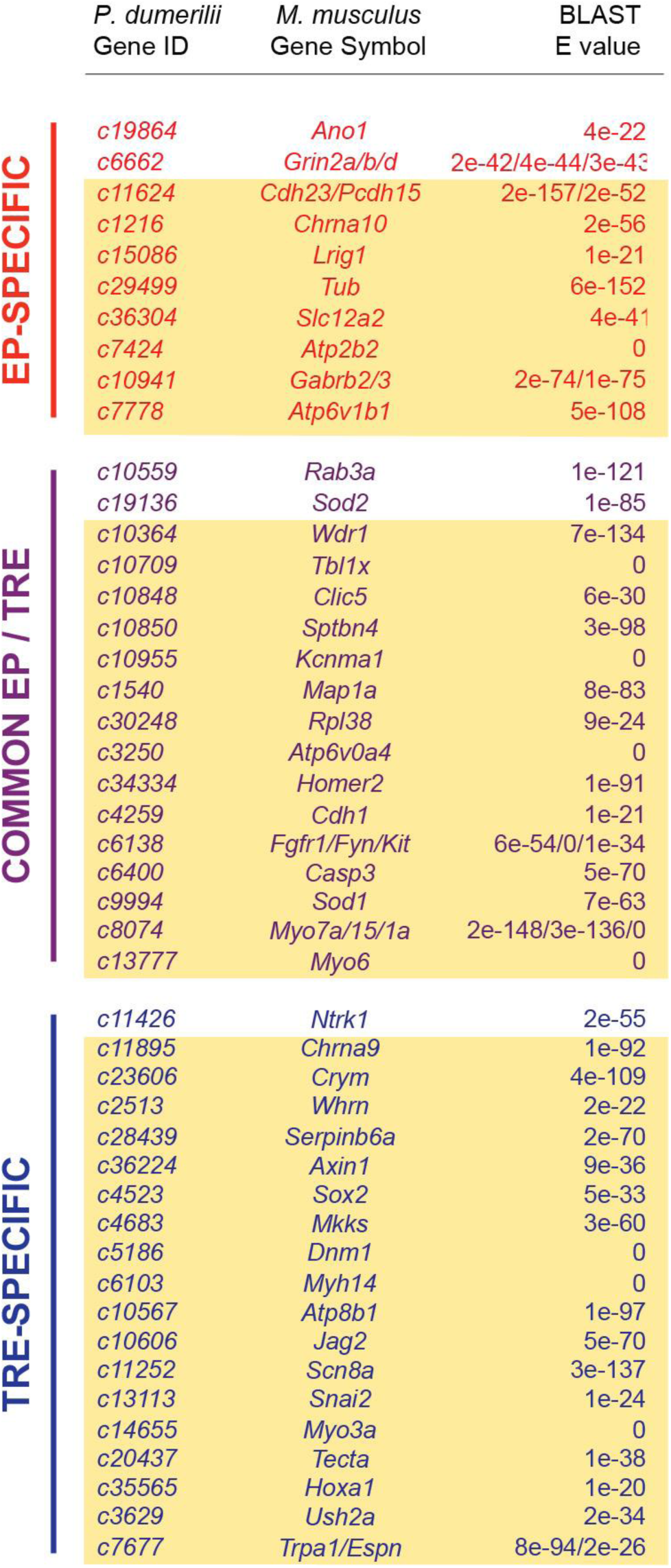
List of the overlapping genes indicated in Fig 3G. Each gene in the “*P. dumerilii* Gene ID” column indicates the best *P. dumerilii* Blast hit to the corresponding *M. musculus* Gene Symbols in the middle column, as described in detail in Methods. The yellow shading indicates genes that are involved in the sensory perception of sound (Fig. 3J).

**Fig. 4-figure supplement 1.**
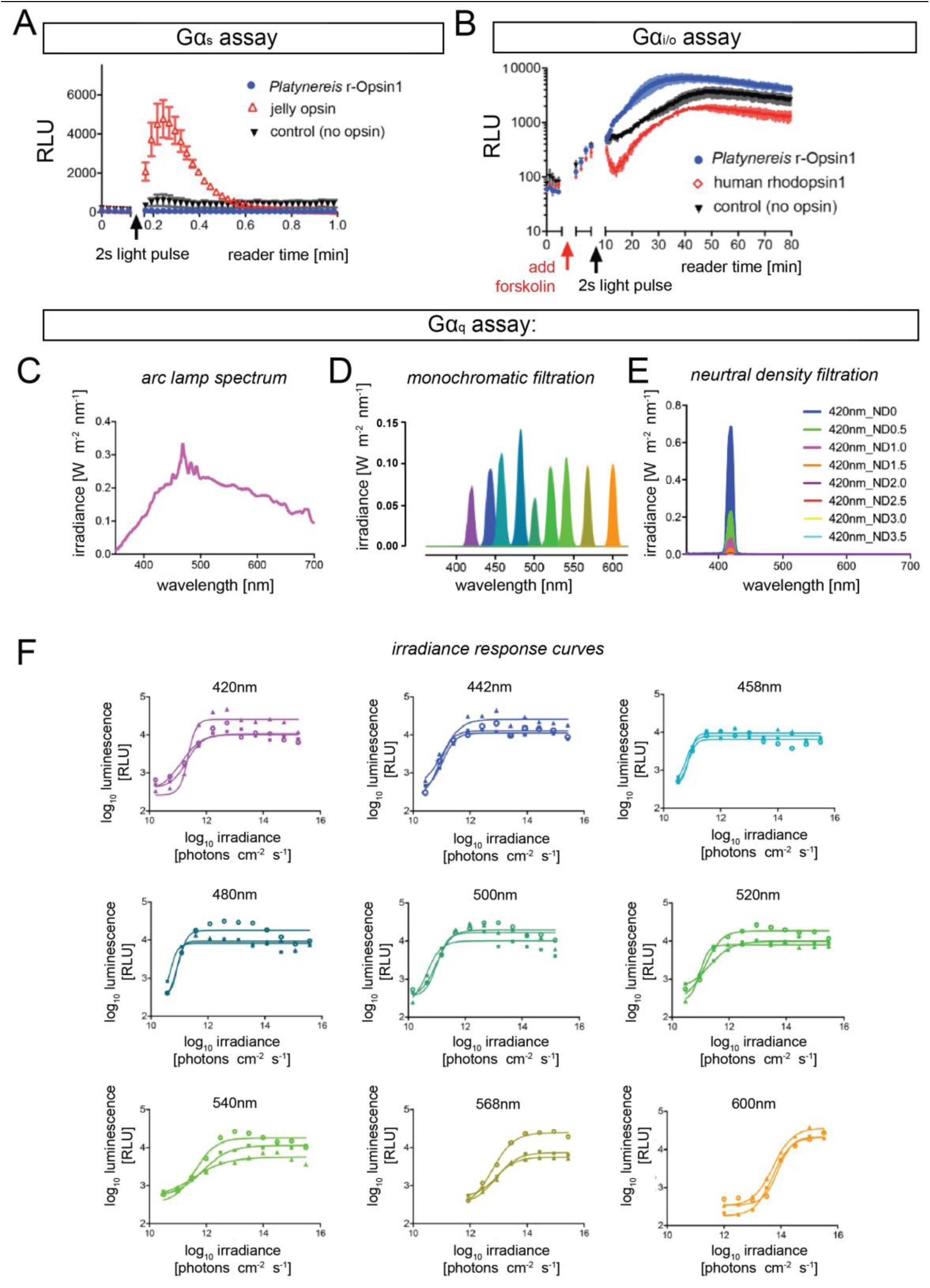
Signaling properties of *Platynereis* r-Opsin1. **(A)** In contrast to reporter cells transfected with a jellyfish *opsin* construct (red; ref. [53]), no detectable luminescence increase is observed after a 30s white light pulse in transfected with *P. dumerilii r-opsin1* (blue), similar to non-transfected controls (black), indicating that *P. dumerilii* r-Opsin1 does not activate Gα_s_. **(B)** Similarly, while human *rhodopsin 1* transfection (red; ref. [53]) makes reporter cells susceptible to a 2s white light pulse (reduction in cAMP concentration in cells pre-exposed to Forskolin), untransfected reporter cells (black) or cells transfected with *P. dumerilii r-opsin1* (blue) do not appear to activate Gα_i/o_ in *P. dumerilii*. In (A) and (B), x axes indicate plate reader time (interrupted by light exposure). Reporters were HEK293 cells transfected with pcDNA5/FRT/TO Glo22F. **(C-F)** Light spectra and irradiance response curves for the Gα_q_ assay presented in **Fig. 4**. (C) Spectrum of the Arc lamp white light used for all G-protein selectivity assays. (D) Monochromatic light produced from the broad-spectrum Arc lamp light using bandpass filters. (E) Example (at 420nm) for the effect of neutral density filters on generating different irradiance levels for test in the irradiance response assays.(F) r-Opsin1 irradiance dose response curves for the Gα_q_ assay shown in **Fig. 4**; panels show the respective wavelength, and the luminescence responses correlated with tested irradiance levels. Respective luminescence values are plotted in relation to the baseline with minimal signal elicited with no light exposure (0% response – no light signal) and the maximum response evoked from that plate (100% response). Each irradiance response curve was fitted with sigmoidal dose response curve to derive the 50% maximal response used to calculate graphs represented in **Fig. 4B,C**. n= 3 independent experiments in all cases.

**Fig. 5- figure supplement 1.**
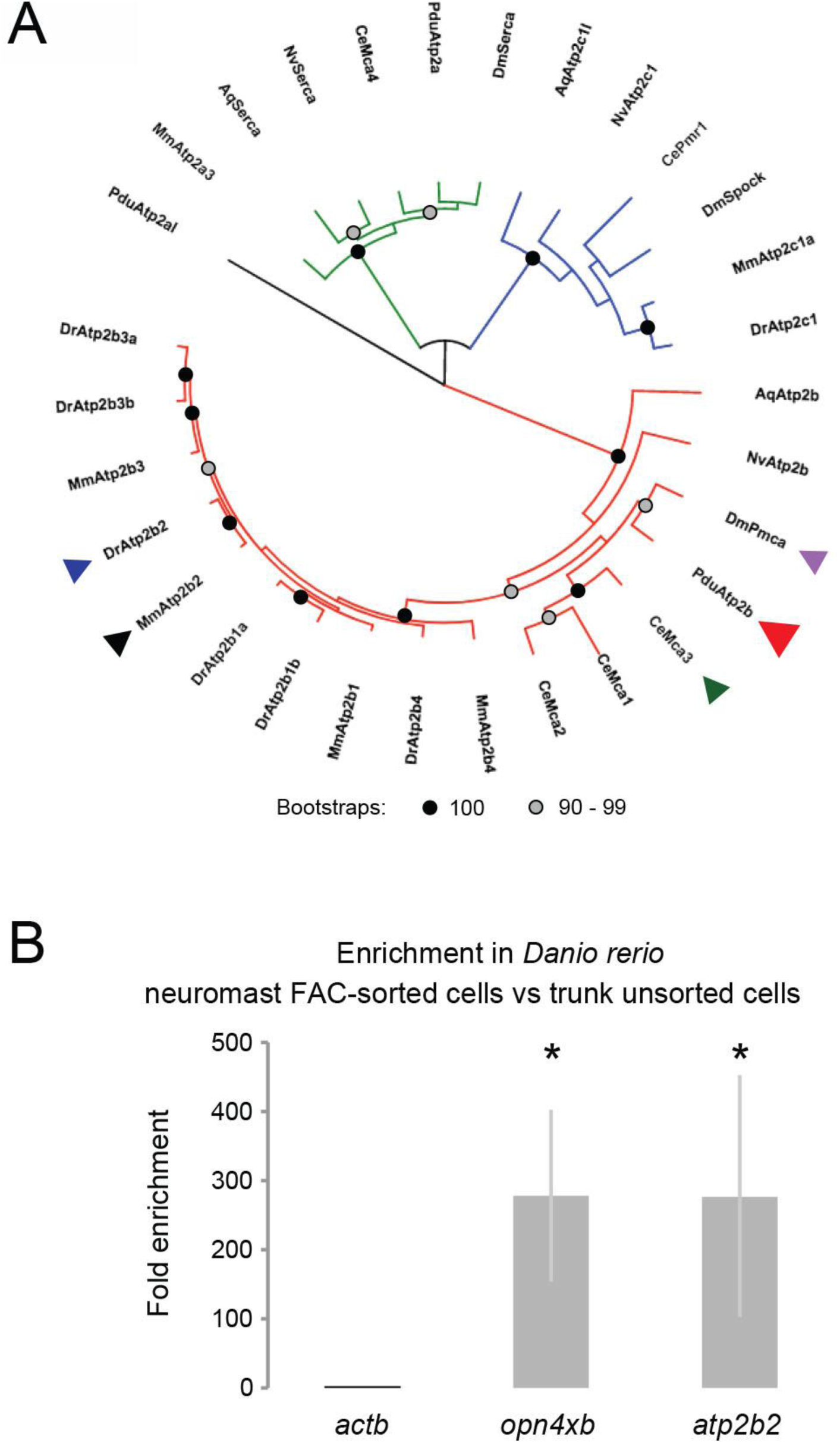
Atp2b2 phylogeny, and enrichment of *atp2b2* in zebrafish neuromasts. **(A)** Phylogenetic tree including Atp2b2 and related genes. Mm: *Mus musculus*; Dr: *Danio rerio*; Dm: *Drosophila melanogaster*; Ce: *Caenorhabditis elegans*; Pdu: *Platynereis dumerilii*; Nv: *Nematostella vectensis*; Aq: *Amphimedon queenslandica*. Red clade: Atp2b protein family; Green clade: Atp2a protein family; Blue clade: Atp2c protein family. Arrowheads indicate proteins encoded by genes mentioned in the manuscript: Black: Mouse Atp2b2; Dark blue: zebrafish Atp2b2; Light blue: Drosophila Pmca; Green: *C. elegans* Mca-3; Red: *Platynereis* Atp2b. **(B)** Bar plots indicate fold enrichment of *actb*, *opn4xb* and *atp2b2* mRNA expression (measured by quantitative PCR) in neuromast cells (FAC-sorted from fish trunks) as compared to unsorted cells from the same fish trunks. Enrichment values were normalized to *actb* levels. As *opn4xb* (previously known as *opn4x2*) has been shown to be specifically expressed in the neuromasts of the lateral line within the trunk of the fish [17], the enrichment of *opn4xb*, confirms the correct isolation of neuromasts cells. * p-value < 0.05 (Student’s t-test).

**Fig. 5- figure supplement 2.**
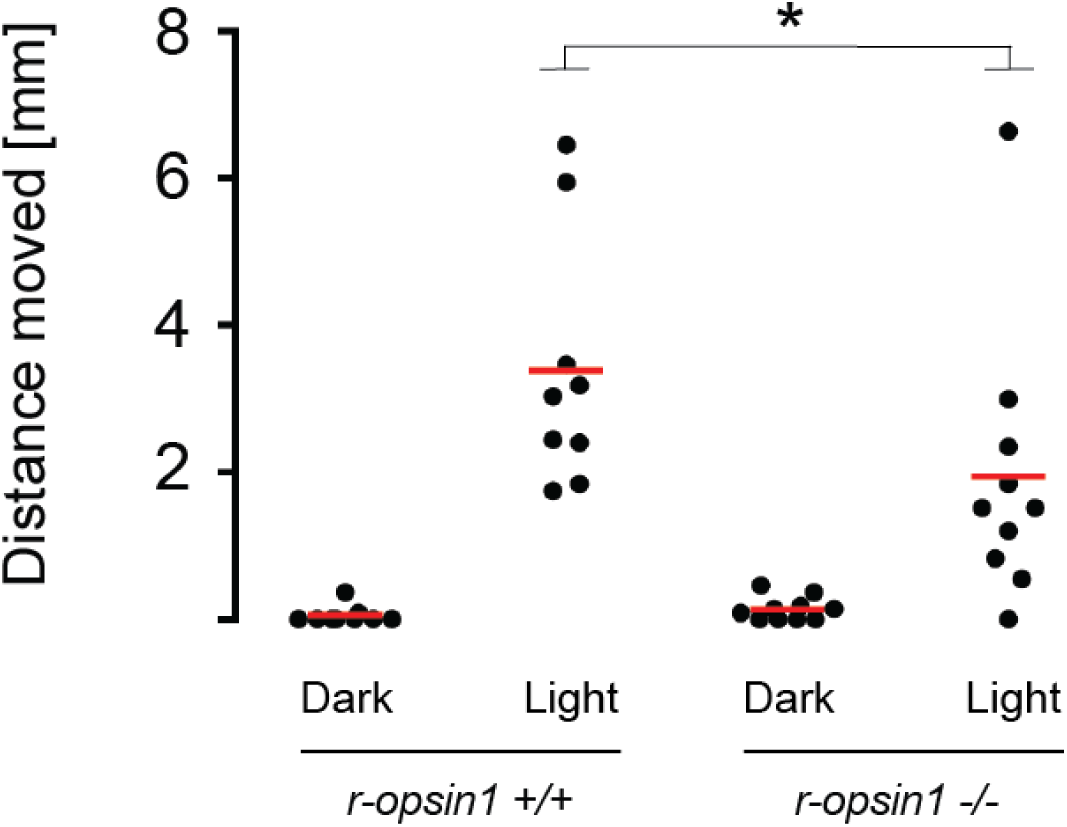
Net avoidance crawling distance of decapitated *r-opsin+/+* and *r- opsin-/-* worms in response to strong light. Dot plot showing the distance moved (in mm) by worms exposed to a bright light pulse (“Light”) or not exposed (“Dark”). * p-value < 0.05 (Wilcoxon rank sum test).

**Fig. 5- figure supplement 3.**
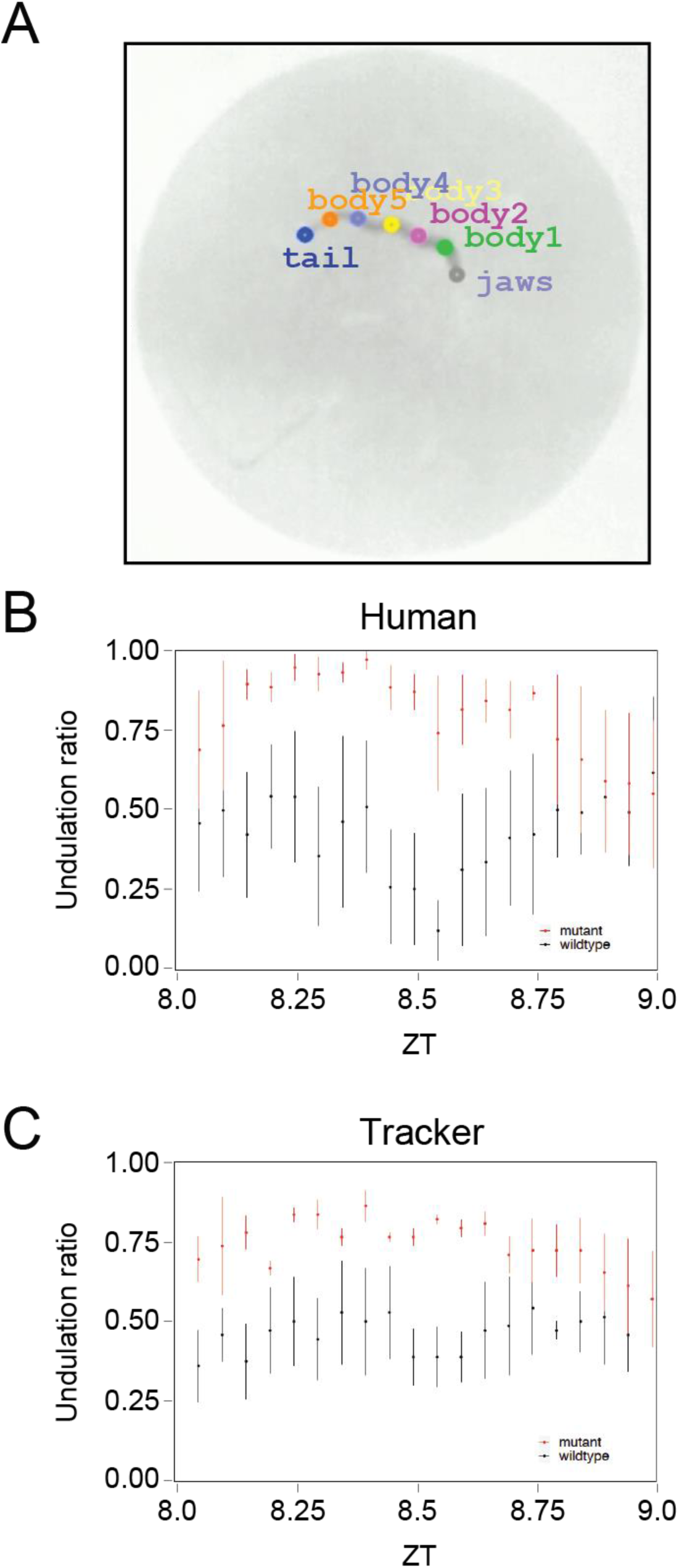
Benchmarking the algorithm used to detect undulation behavior. **(A)** Behavior arena of an individual worm showing the key points detected by the automatized tracker along the worm: jaws (“jaws”), 5 points along the trunk (“body1-5”) and tail (“tail”). **(B,C)** Undulation ratio of wild-type (black) and mutant (red) worms as determined by a human observer (B) or by the automatized tracker (C). Each point represents the mean of all wild-type or mutant worms within a 3-minute window, and vertical bars represent the standard error of the mean (n=4 for each genotype).

## METHODS

### Animal culture and handling

All animal research and husbandry was conducted according to Austrian and European guidelines for animal research (fish maintenance and care approved under: BMWFW-66.006/0012-WF/II/3b/2014, experiments approved under: BMWFW-66.006/0003-WF/V/3b/2016, which is cross-checked by: Geschäftsstelle der Kommission für Tierversuchsangelegenheiten gemäß § 36 TVG 2012 p. A. Veterinärmedizinische Universität Wien, A-1210 Wien, Veterinärplatz 1, Austria, before being issued by the BMWFW). Zebrafish were kept in a constant recirculating system at 26-28°C in a 16h light / 8h dark cycle. Collected embryos were kept at 28°C until hatching.

*Platynereis dumerilii* were raised and bred in the Max Perutz Labs marine facility according to established procedures [76]. Experimental animals were immature adults fed last 4 to 6 days prior to the day of the experiment. Remaining food was removed a day after feeding, and the seawater changed, leaving the worms unperturbed for 3 to 5 days prior to sampling. All pMos{rops::egfp}^vbci2^ transgenic worms [17] used for transcriptome profiling were screened for strong EGFP fluorescence under a stereo microscope system (Zeiss SteREO Lumar V12) at least 6 days before the experiment. To partially immobilize the worms for the screening, worms were shortly transferred to a dry petri dish.

### Fluorescence-Activated Cell (FAC) Sorting

EGFP-positive cells from 1–2 worms were isolated by FAC sorting with three biological replicates. For the head and trunk of each replica, a sample of unsorted cells was also isolated as reference.

To FAC-sort EGFP+ cells, 1–2 immature transgenic worms per biological replicate were decapitated under a stereoscopic microscope (Zeiss Stemi 2000; Zeiss, Germany) by using a sterile scalpel (Schreiber Instrumente #22; Schreiber Instrumente GmbH, Germany). Separated heads or trunks were placed on ice for about 2 min in 2ml seawater immediately before dissociation. Heads were mechanically dissociated through a nylon 70µm cell-strainer (Falcon, USA) in 600µl seawater. Trunks were first cut into 3-4 pieces using a sterile scalpel, and then dissociated in the same way, using 3ml seawater. Cell suspensions were passed four times through 35µm nylon mesh cell-strainers (5ml Polystyrene round-bottom tube with cell-strainer cap, Art. #352235, Falcon), and placed on ice. Finally, the volume of the single-cell suspensions was adjusted to 600µl (for heads) or 3ml (for trunks) with ice-cold seawater. Heads and trunks from 1-2 non-transgenic worms were also dissociated as negative controls for the detection of EGFP fluorescence.

Cell suspensions were stained with Propidium Iodide (PI; ThermoFisher Scientific, P1304MP) by adding 8µl of 1.5mg/ml PI per ml of cell suspension, and were kept on ice until FAC-sorted. Stained cell suspensions were analyzed on a FACSAria IIIu FAC Sorter (BD Bio-sciences). FAC-sorting (FACS) events were first gated to exclude aggregates using the FSC-A and FSC-W channels. To separate real EGFP fluorescence from autofluorescence, we followed a previously established strategy [77], measuring fluorescence elicited by a 488nm laser using two distinct detectors (see **Fig. 1C,D**). One quantified fluorescence in the 515-545nm range (“FITC” axis in **Fig. 1C,D**; **Fig.1-figure supplement 1A,B**), while the other quantified fluorescence in the 600-620nm range (“PE” axis in **Fig. 1C,D**; **Fig.1-figure supplement 1A,B**). Comparison between stained cell suspensions from transgenic (**Fig. 1C,D**) and wild-type (**Fig.1-figure supplement 1A,B**) specimens allowed for the definition of the gate containing EGFP+ events (boxes in **Fig. 1C,D**).

### Transcriptome profiling of EGFP+ cells

Aliquots of 30 – 120 FACS events from the EGFP+ gate of transgenic heads or trunks were sorted into wells of a 96-well plate (Hard-Shell Low-Profile Thin-Wall 96-Well skirted PCR plate, Bio-Rad HSP-9631) containing 4µl of lysis buffer. The lysis buffer consisted of 3.8µl of 0.2% (vol/vol) Triton X-100 (20μl Triton X-100 BioXtra, Sigma T9284 in 10ml Nuclease-free H2O) + 0.2µl RNase Inhibitor (Clontech 2313A). Loading of the plate was carried out under a laminar flow hood to avoid contamination, and according to the recommended procedures for subsequent isolation of RNA and synthesis of cDNA using the Smart-Seq2 technology [25]. The 96-well plate containing lysis buffer was kept on ice until loaded onto the FAC-sorting machine. The 96-well plate was maintained at 4°C during the FAC-sorting procedure.

A sample of unsorted cells was also taken from the same cell suspension from which the FAC-sorted cells were isolated. For this, immediately prior to FAC-sorting, 0.4 μl of the cell suspension was pipetted into 4μl lysis buffer onto the same 96-well plate used for the sorted cells. From then on, the lysates with FAC-sorted cells and the lysates with unsorted cells were subjected to the same procedures. Immediately after sorting, the 96-well plate containing the lysates was sealed (AlumaSeal CS Films for cold storage, Sigma-Aldrich Z722642-50EA) and stored at -80°C.

### Bioinformatic Analyses

#### Transcriptome assembly

All sequencing reads from head or trunk FAC-sorted and unsorted samples from transgenic worms were used to assemble a *de novo Platynereis dumerilii* transcriptome, using the Trinity Software version 2.0.6 [78]. Transcripts were filtered for a minimum length of 250 bp. Also, all transcripts that contained overlapping sequences of 50 bp or longer were grouped into clusters. This ensured that each sequencing read (50bp) could be unambiguously mapped onto a single cluster. For each cluster, we computed nominal transcript length by concatenating the unique sequences within the cluster.

#### Mapping reads to transcriptome

Sequencing reads from each individual sample were mapped onto the de novo transcriptome using the NextGenMap program [79]. Reads that could be mapped onto more than one transcript within the same cluster were mapped only onto one of the transcripts. The number of reads mapped onto each transcript were counted, and counts onto the transcripts within each cluster were added to obtain the total number of reads per cluster. As different transcripts within each cluster likely reflect polymorphisms and splice variants, we refer to these clusters as “genes”. Gene contigs corresponding to spiked-in sequences (obtained by blast against ERCC92 sequences) were removed to obtain the list of *P. dumerilii* genes.

#### Determining gene expression levels

To obtain normalized expression levels for each gene, we computed the number of transcripts per million reads (TPMs) as follows: 1. We assigned a nominal transcript length to each gene by concatenating the longest transcript within the cluster with all the non-overlapping sequences of the rest of the transcripts of the cluster; 2. For each gene, we normalized the read counts to the associated transcript length (in kilo base pairs); 3. For each sample, we normalized to the total million reads in the sample. Genes were considered to be expressed in any given sample if they showed >= 12 TPMs in at least one biological replicate. This threshold is consistent with our enrichment analysis (see below), since it is approximately the minimum expression level required for a gene to be significantly enriched in our differential expression analysis. A gene was considered to be expressed specifically in EP (or TRE) cells if it was expressed in EP (or TRE) cells, and not in TRE (or EP) cells.

#### Differentially expressed genes

To identify differentially expressed genes, we used the EdgeR software package, according to the developerś instructions [27, 80]. For each experiment, we used the raw read counts to first filter out all genes that did not have more than 1 count per million in at least 3 samples within the experiment, and to then calculate normalization factors for each sample by comparing all samples of the same experiment. Subsequently, we used the quantile-adjusted conditional maximum likelihood (qCML) method to calculate the common and gene-wise dispersion, and the exact test for the negative binomial distribution to test for differentially expressed genes [27, 80]. Only genes with an FDR <= 0.05 were considered significantly differentially expressed. Genes were considered significantly enriched in EP (or TRE) cells of the head (or the trunk) if they fulfilled the following two criteria: (i) They were identified as differentially expressed between EP (or TRE) cells of the head (or the trunk) and unsorted cells of both head and trunk; (ii) their expression in EP (or TRE) cells of the head (or the trunk) was higher than in unsorted cells of the head and the trunk. Genes were considered specifically enriched in the EP (or TRE) cells of the head (or the trunk), if they were enriched in the EP (or TRE) cells of the head (or the trunk), and not enriched in the TRE (or EP) cells of the head (or the trunk). Two genes (c8629 and c14134) were excluded from further analyses because they represent redundant fragments of the *ropsin1* gene.

#### Detection of *bona fide* homologs of *Drosophila* and mouse genes

To systematically assess putative gene homology relationships between *Drosophila melanogaster* or *Mus musculus* and *Platynereis dumerilii*, we used the *tblastn* algorithm to compare all *Drosophila melanogaster* or *Mus musculus* protein sequences in the ENSEMBL database (Drosophila_melanogaster.BDGP6.pep.all.fa (10th March 2016); Mus_musculus.GRCm38.pep.all.fa (10th March 2016)) to all transcripts in our *P. dumerilii de novo* assembled transcriptome. To each *D. melanogaster* or *M. musculus* gene ID, we assigned the *P. dumerilii* gene with the best *tblastn* hit, with a stringent E value threshold of 1E-20.

#### Identification of *P. dumerilii* components of the phototransduction pathway

To identify *P. dumerilii* components of the canonical r-Opsin phototransduction pathway, we assigned *bona fide P. dumerilii* homologs to the key components of the *D. melanogaster phototransduction pathway* (**Fig. 2B**; R-Opsin, Gaq, Gb, Gg, NorpA/PLC, INAC/PKC, Trp, Trpl, INAD, Cam, NINAC/MyoIII; Arrestin2, PIP5K). The corresponding ENSEMBL (Drosophila_melanogaster.BDGP6.pep.all.fa (10th March 2016)) gene symbols are as follows: R-Opsin: NinaE/Rh3/Rh4/Rh5/Rh6; Gaq: Galphaq; Gb: Gbeta76C; Gg: Ggamma30A; NorpA/PLC: NorpA; INAC/PKC: InaC; Trp: Trp; Trpl: Trpl; INAD: InaD; CaM: Cam; NINAC/MyoIII: NinaC; Arrestin2: Arr2; PIP5K: PIP5K59B. To each *D. melanogaster* protein we assigned the best *P. dumerilii tblastn* hit, with an E value threshold of 1e-20, as described above. Two proteins (Gg and InaD) had no *P. dumerilii tblastn* hits that satisfied this stringent threshold. Therefore, to assign *P. dumerilii* homologs to these proteins, we lowered the stringency of the E value threshold to 1e-8. To corroborate that *c33855* is a *bona fide* homolog of Gg (E value 2e-9), we confirmed that this *P. dumerilii* gene is the best *tblastn* hit of the *M. musculus* Gg counterpart (Gng; E value against *c33855*: 2e-9). Similarly, to corroborate that *c7982* is a homolog of InaD (E value 2e-16), we confirmed that this gene is the best *tblastn* hit of the *M. musculus* InaD counterpart (Mpdz; E value against *c7982*: 2e-81).

#### Statistical assessment of subset specificity

To assess whether the number of EP- and/or TRE-expressed/enriched genes overlapping with the *P. dumerilii* homologs of a set of N *D. melanogaster* or *M. musculus* genes was meaningful, we generated 10^4^ sets of N randomly-picked *D. melanogaster* or *M. musculus* genes, and performed the same analysis as for our real set of N *D. melanogaster* or *M. musculus* genes. We then determined the frequency with which such randomly generated sets resulted in an overlap that was equal or higher than that found for our real set.

#### Molecular phylogenetic analysis of plasma membrane Calcium-Transporting ATPases and related proteins

Candidate Plasma membrane Calcium-Transporting Atpases and related proteins were identified from the *Platynereis dumerilii* transcriptome with the *tblastn* algorithm, using selected animal homologs as query (see **Fig.5-figure supplement 1A**). Predicted sponge proteins were aligned with their counterparts from other animals using MUSCLE [81], and molecular phylogenetic analyses were performed using the IQTREE software [82].

### Analysis and validation of differentially expressed genes

To validate the results of our differential expression analysis, we selected 2 common EP-/TRE-enriched genes (*ngbl/c10609* and *tmdc/c2433*), and 3 TRE-specific genes (*f8a/c6996*, *dmd/c7924* and *trpA/c7677*). The genes selected cover a wide FDR range in our statistical analysis (**Fig.2-figure supplement 2**). *ngbl/c10609* and *tmdc/c2433* are among the top enriched genes in both EP and TRE samples (FDR < 0.01), whereas *trpA/c7677* (FDR = 0.038) is close to the significance threshold (**Fig.2-figure supplement 2**). The low FDR values for *ngbl/c10609* and *tmdc/c2433* reflect the high level of expression of these genes in the EP and TRE samples for all three biological replicates, and the low level of expression in the unsorted samples (**Fig.2-figure supplement 1A,B**). From these data, we expected that *ngbl/c10609* and *tmdc/c2433* would be expressed at low levels (or not expressed at all) in any cell type other than EP and TRE cells. We used the established single- or two-color whole mount *in situ* hybridization (WMISH) [75] with *r-opsin1* as reference. Within the head, *r-opsin1* is prominently expressed in the four adult eyes [17], which is reproduced in our controls (**Fig.2-figure supplement 3C,D**, detected in red). Of note, a dense pigment cup covers the internal portion of each eye that contains the photosensitive outer segments of the retinal photoreceptors [31]. This pigmented area can be seen as a dark area in the eyes (**Fig.2-figure supplement 3C.D**), which partially shields the *r-opsin1* staining. However, due to the localization of the photoreceptor cell bodies (and those of the support cells) outside the pigment cup, gene expression can be assessed in this apparent circle around the pigment cup (broken white contour in **Fig.2-figure supplement 3D**). In this non-pigmented area of the eyes, the red staining for *r-opsin* was clearly discernible (**Fig.2-figure supplement 3D**). Single-color ISH using a probe against *ngbl*/*c10609* showed expression of this gene in the EP as well (**Fig.2-figure supplement 3E,F,** blue staining), confirmed by two-color-WMISH (**Fig.2-figure supplement 3G,H,** arrowhead).

The TRE cells in the trunk of the worm are apparent as single, *r-opsin1-*positive cells within each parapodium in the ventral flap of the dorsal parapodial arm (ref. [17]; **Fig. 3D,E** and **Fig.2-figure supplement 4A,D,E**, red staining). When tested on trunk samples, the probe for *ngbl*/*c10609* revealed a similar expression pattern to *r-opsin1* (**Fig.2-figure supplement 4B**, blue staining). Two-color-WMISH confirmed the co-expression (**Fig.2-figure supplement 4C**, purple color, arrowhead).

Similarly, a riboprobe against *tmdc/c2433* revealed specific staining in the EP cells as well as single cells within each parapodium, in a position consistent with the TRE cells [32].

Specific expression of the three selected TRE-specific genes, was also validated using single- and double-WMISH. A probe against *trpA*/*c7677* shows no expression in the eyes (**Fig.2-figure supplement 3I,J**), while *trpA/c7677* is detected in small spots in each parapodium (**Fig.2-figure supplement 4F,G**, blue staining). Two-colour-WMISH shows that one of the spots in each parapodium overlaps with *ropsin1* expression, limited to only part of the cell (**Fig. 3E**). *f8a*/*c6996* and *dmd/c7924* were also expressed in a single cell in the ventral flap of the dorsal arm of each parapodium, in a position that is consistent with the TRE cell (**Fig.2-figure supplement 4H-K**), while expression in the eyes was undetected (*f8a/c6996*; **Fig.2-figure supplement 3K,L**) or extremely weak (*dmd/c7924*; **Fig.2-figure supplement 3M,N**). Along with our set of control genes, these additional validations yield a total of 10 genes that confirm our enrichment analysis (**Fig.2-figure supplement 2**). The confirmation of *f8a/c6996*, *dmd/*c*7924* and *trpA/c7677*, with relatively low level of expression and moderate enrichment FDR values (**Fig.2-figure supplement 2**) particularly strengthens the validity of our analysis. It is worth noting that genes expressed at low levels are more likely affected by stochasticity effects during cDNA synthesis and amplification than their highly abundant counterparts. This provides a likely explanation why *dmd*/*c7924* and *trpA*/*c7677* are detected, respectively, in two and one of the three biological replicates in TRE cells (**Fig.2-figure supplement 1D,E**).

### Bioluminescence assays to assess Gαq, G*α*s and G*α*i/o coupling

To test whether *P. dumerilii* rOpsin1 can activate G*α*q, G*α*s or G*α*i/o GPCR signaling upon light exposure, we adapted established cell culture second messenger assays [51]. For this, the *P. dumerilii r-opsin1* gene was heterologously expressed in HEK293 cells. Co-transfected luminescence reporters assessed either activation of G*α*q signaling (pcDNA5/FRT/TO mtAeq; expressing *aequorin* as reporter of intracellular calcium [83]) or activation of G*α*s or G*α*i/o signaling (pcDNA5/FRT/TO Glo22F). Transfected cells were incubated overnight with the chromophore 9-cis retinal in single wells of a 96-well plate. Cells were then incubated with 10µM Coelenterazine (for G*α*q) or 0.1M Luciferin (for G*α*s or G*α*i/o, respectively) in the dark for 2 hours, and were subsequently exposed to a 2s (for G*α*q and G*α*i/o) or 30s (for G*α*s) pulse of white light. The white light pulse was generated by an Arc lamp, spectrum in **Fig.4-figure supplement 1C**. Raw luminescence was measured from each single well on a Fluostar Optima plate reader (BMG Labtech, Germany). While the well under recording was exposed to the light pulse, all other wells were protected from light with a black sheet. To assess activation of G*α*q signaling, luminescence was measured with a resolution of 0.5s and cycle of 2s. To assess activation of G*α*s signaling, increase in cyclic Adenosine Monophosphate (cAMP) levels was assessed by measuring luminescence with a resolution of 1s and cycle of 30s. To assess activation of G*α*i/o signaling, cells were treated with 2µM Forskolin (Sigma-Aldrich) prior to the light pulse, and decrease in cAMP levels was assessed by measuring luminescence with a resolution of 1s and cycle of 30s. The time between the light exposure to the well and the recording of raw luminescence measurements was approximately 3s. Measurements taken during the dark incubation preceding the light pulse were used as baseline. As positive controls, we used established constructs for jellyfish Opsin (G*α*s assay; ref. [53]), human Rhodopsin 1 (G*α*i/o assay; ref. [53]) and human Opn4 (G*α*q assay, ref. [51]).

#### Measurement of spectral sensitivity of *P. dumerilii* r-Opsin1

To determine the spectral sensitivity of *Platynereis* r-Opsin1, the aforementioned bioluminescence assay was further refined. Band-pass (420, 442, 458, 480, 500, 520, 540, 568 and 600nm) and neutral density filters (0-3.5) (**Fig.4-figure supplement 1D,E**) were used to deliver defined irradiance doses of distinct wavelengths in 2s light pulses. Maximum luminescence levels acquired from three independent replicates were plotted against the respective irradiance doses used. Individual irradiance response curves for a given wavelength were then fitted to a sigmoidal dose response function (variable slope, minimal asymptote value constrained to the average raw luminescence baseline for each wavelength), allowing to derive EC_50_ values (irradiance required to elicit half-maximal luminescence responses) for each wavelength (**Fig.4-figure supplement 1F**). The relative sensitivity at each wavelength was calculated as described in ref. [51]. Likewise, the fitting of data to the Govardovskii visual templates for each wavelength, and the determination of the curve with the best fit to the measured data to determine λ_max_ of *Platynereis* r-Opsin1 in cell culture (**Fig. 4B,C**) followed established procedures [51].

### Enrichment of *atp2b2* mRNA expression in zebrafish neuromasts cells

The *TG(pou4f3:GAP-GFP)^s356t^* transgenic zebrafish line was used for this experiment, which expresses membrane-targeted GFP under the control of the *brn3c* promoter/enhancer [49]. 30 transgenic or non-trangenic larvae at 6-10 days post-fertilization were decapitated under a stereoscopic microscope. Trunks were dissociated by incubating them in 0.5% Trypsin-EDTA 10x (59418C-100ml Sigma Aldrich) diluted in PBS for 3-4 min, and shearing through a 1ml pipette tip for an additional 6 min. Cell preparations were filtered once through a 70 µm cell-strainer (Falcon, USA), and three times through 35µm nylon mesh cell-strainers (5ml polystyrene round-bottom tube with cell-strainer cap, Art. #352235, Falcon), and were then placed on ice.

Cell suspensions were stained with propidium iodide (PI; ThermoFisher Scientific, P1304MP) by adding 8µl of 1.5mg/ml PI per ml of cell suspension, and were kept on ice until FAC-sorted. To isolate GFP^+^ neuromast cells cell suspensions were analyzed on a FACSAria IIIu FAC Sorter (BD Bio-sciences), using the same gating strategy as above for the isolation of EGFP^+^ cells from *Platynereis*. Non-transgenic cell preparations were used to distinguish EGFP^+^ cells from autofluorescent cells, and therefore be able to accurately design the EGFP^+^ gate. EGFP^+^ cells were directly FAC-sorted into RLT lysis buffer (Qiagen). After collection, Lysate was vortex for 30 sec and stored at -80°C. A sample of unsorted cell preparation was lysed, to be used as unsorted sample.

Total RNA was isolated from EGFP^+^ FAC-sorted and unsorted cell lysates by using the RNeasy mini kit (Qiagen) according to manufacturer’s guidelines, cDNA was synthesized by using the QuantiTect Reverse Transcription kit (Qiagen) according to manufacturer’s guidelines. To measure gene expression levels of *actb, opn4xb* and *atp2b2*, quantitative PCR (qPCR) was performed on 96-well plates, in a StepOne Real-Time PCR System (Applied Biosystems) using SybrGreen chemistry (Thermo Fischer Scientific). The total volume of all qPCR reactions was 20 µl. Measured expression levels were used to calculate enrichments, normalizing to the *actb* levels. Statistical significance of enrichment was tested on the QPCR relative number of cycles at threshold (cycles at threshold for *opn4xb* or *atp2b2* relative to *actb*) in EGFP+ samples compared to unsorted samples. Bartlett’s test was used to test for equal variance.

### Behavioral analyses

#### Light-induced crawling movement

To assess the light-induced crawling response of immature wild-type and *r-opsin1* mutant trunks, we followed a previously established method [17]. For both wild-type and *r-opsin1* mutant genotypes, we used the pMos{rops::egfp}^vbci2^ transgenic background. Animals were screened prior to the assay to ensure similar EGFP fluorescence intensity.

#### Undulation Behavior Analysis

Wild-type and *r-opsin1* mutant genotypes were used in the pMos{rops::egfp}^vbci2^ transgenic background. Worms were kept unfed for 3 days prior to the start of the experiment. On the day of the start of the experiment, worms were decapitated and then placed in individual hemispherical concave wells of a custom-made 25-well clear plate [30, 63]. To obtain trunks, specimens were anesthetized by using a 1:1 solution of seawater and 7.5% MgCl_2_, placed on a microscope slide under a binocular dissecting microscope, and decapitated using a surgical blade (#22; Schreiber Instrumente GmbH, Germany). To increase the chance that decapitated worms could build tubes, the decapitation plane was chosen anterior to the pharyngeal region.

Video recording of worm behavior over several days was accomplished as described previously [59, 63]. Prior to recording, worms were incubated for 2– 4 hours to allow them to build tubes, which is part of their normal behavior. During the recording, worms were subjected to one complete light-dark cycle (16 h light/8 h darkness), followed by 4 days of constant darkness. White light was generated by custom made LEDs (Marine Breeding Systems, St. Gallen, Switzerland), reaching worms with an intensity of 5.2×10^14^ photons/cm^2^/s. Analyses focused on Zeitgeber Time (ZT) 6-14 of the LD cycle (LD1), and circadian time (CT) 6-14 of the first DD cycle (DD1). ZT0: start of lights on. Worms that had not built a tube during the first hours of the recording, or those that had matured by the end of the experiment, were excluded from further analysis.

Undulation analysis was performed using positional data of 7 discrete body points (**Fig.5-figure supplement 3A**), obtained via a deep learning based key point prediction algorithm. The algorithm/neural network was created via the interface of Loopy, developed by loopbio GmbH (Vienna, Austria, http://loopbio.com). For training the network, points were manually annotated using 2,740 individual frames obtained from different recordings with the set-up described above. To ensure high diversity of the training set, chosen recordings covered different sizes and shapes of worms as well as different times of the day. The subsequent data analysis was carried out in Python 3.7.9 using the SciPy (1.5.2), pandas (1.1.3) and NumPy (1.19.2) packages [84–86].

The positional data was first checked for sufficient prediction coverage: worms for which any single point was annotated in less than 90% of the frames were excluded from further analysis. For the retained individuals, any missing XY values were inferred linearly from non-missing data. To identify undulation, power spectral density was estimated on 10 second intervals for the position of each body point excluding the jaw and the tail by means of a periodogram. For every point, the dominant frequency within the given time-window was determined. A movement was defined as undulation if any of the 5 body points showed a total movement of 0.5 – 10 pixels and had a dominant frequency within a range of 0.5 – 1.5Hz. Undulation ratios obtained by manually scoring video segments were used to benchmark the automated algorithm (**Fig.5-figure supplement 3B,C**).

All statistical tests were done using R (version 3.6.1). First, from the undulation ratios the area under the curve was calculated for every replicate and then the datasets were tested for normal distribution (Shapiro-Wilk normality test). To determine if there were differences between the groups, either a paired (light versus dark) Wilcoxon signed rank test or an unpaired (wildtype versus mutant) Wilcoxon rank sum test was conducted. Results were considered statistically significant with a p-value <0.05.

### Transcriptome profiling

After addition of 2µl of dNTP mix (10mM each; Fermentas, R0192), 2µl of oligo-dT-30VN primer (10µM; 5’ – AAGCAGTGGTATCAACGCAGAGTACT30VN-3’), and ERCC spike-in RNA (Ambion) (1:1,000,000 dilution) to the lysates of FAC-sorted or unsorted cells, mRNA isolation, cDNA synthesis with amplification was performed according to the standard Smart-Seq2 protocol [25]. Single-end 50bp-read sequencing of the cDNA libraries was performed on an Illumina HiSeq3000/4000 platform according to the manufacturer’s protocol. For all samples, transcriptome profiles for three independent biological replicates were obtained.

### *In-situ* hybridization and imaging

In-situ hybridization and dual-color in-situ hybridization on whole heads and trunk pieces (5-10 segments) of immature worms were performed according to established methods [17]. Whole heads and trunk pieces were mounted on glass slides and imaged on a Zeiss Axio Imager with 10x or 40x oil immersion objectives. Single parapodia were cut out of the trunk pieces, mounted on glass slides and imaged with 10x or 40x oil immersion objectives. A Zeiss Axiocam MR5 camera was used for documentation of stainings.

### Generation of *r-opsin1* mutant strains

The *r-opsin1* genomic region was amplified to screen putative size polymorphic alleles or single nucleotide polymorphisms (SNPs) from different *Platynereis* strains (PIN, VIO and ORA) using the following primer combinations: rops1_F1/R1, rops1_F2/R2, rops1_F3/R3, rops1_F4/R4 and rops1_F5/R5. The target alleles or SNPs were screened as described in ref. [20]. *r-opsin1* TAL Effector Nuclease (TALEN) pairs were designed in several non-polymorphic exon regions using the TALE-NT prediction tool [87]. *In silico* predictions were performed by using customized design conditions, 15 left/right Repeat Variable Diresidue (RVD) length, 15-25bp spacer length, G substitute by NN RVD and presence of exclusive restriction enzyme site around the spacer region. The predicted *r-opsin1* TALENs were constructed *in vitro* using Golden Gate assembly protocol (Golden Gate TAL Effector Kit 2.0, Addgene #1000000024) [88]. The final TALEN repeats were cloned to heterodimeric FokI expression plasmids pCS2TAL3-DD for left TALEN array and pCS2TAL3-RR for right TALEN array [89]. All cloned TALEN plasmids were sequence-verified using TAL_F1 and TAL_R2 primers. *r-opsin1* TALEN mRNA for each array were made by linearizing the corresponding plasmid by NotI digestion and transcribed *in vitro* using mMESSAGE mMACHINE Sp6 kit. Two TALEN pairs targeting exon 1 of *r-opsin1* were designed and generated using the above *in vitro* assembly protocol. Both *r-opsin1* TALEN spacer regions were flanked with restriction sites (TAL 1 – Bts1 and TAL 2 – Taa1). Following microinjection of 200ng/µl *r-opsin1* TALEN mRNA, the *Platynereis* embryos were screened for mutations using incomplete restriction digestion and confirmed by sequencing the undigested band. Several injected embryos were raised and outcrossed to wildtype. The F1 outcrossed worms were screened for mutations with a similar restriction digest procedure. Two deletion and insertion mutations were recovered (17bp deletion and 1bp deletion). Mutant worms were raised and crossed for several generations to generate both homozygous incross strains and respective wild-type relatives.

### Light and temperature conditions

*r-opsin1*-mutant pMos{rops::egfp}^vbci2^ worms and the corresponding pMos{rops::egfp}^vbci2^ control individuals used for transcriptomic analysis, were incubated without feeding for 3 - 5 days before the experiment. Blue light of 470nm was generated using LEDs. The resulting spectrum and intensity of the light was measured using a SpectriLight ILT950 Spectroradiometer (International Light Technologies, MA, USA) (**Fig. 5B**). The temperature (kept between 18.5 and 20°C) was monitored during the 3 - 5 days of blue light incubation using a HOBO Pendant Temperature/Light Data Logger (Part #UA-002-64, Onset Computer Corporation, MA, USA).

EGFP transgenic worms used for transcriptomic analysis at distinct light conditions were incubated for 3 - 5 days in blue light or in dim white light after decapitation. Blue light conditions were as described above. Dim white light conditions were obtained by placing the worms in an area partially protected from light within a room with standard white light illumination. The exact spectrum and intensity of the light (see **Fig. 5B**) was determined using the same spectroradiometer as described above. The temperature was monitored with a similar device as described above, and was kept within the same range as in the blue light conditions (18.5 to 20°C).

### Metadata/ source files availability

All metadata and source files are available for download from DRYAD: https://datadryad.org/stash/share/_AWaPRHfmuHKyq9CzHt7YNtmaAgHFQXjfklkVE1n6SI.

doi:10.5061/dryad.m63xsj416

This includes raw data, scripts, and the newly assembled and size-filtered transcriptome, used for quantitative mapping (cf. section on Transcriptome profiling).

## REFERENCES

[1] Arendt D, Wittbrodt J. Reconstructing the eyes of Urbilateria. Philos Trans R Soc Lond B Biol Sci 2001;356:1545–63. doi:10.1098/rstb.2001.0971.

[2] Arendt D, Tessmar K, de Campos-Baptista M-IM, Dorresteijn A, Wittbrodt J. Development of pigment-cup eyes in the polychaete *Platynereis dumerilii* and evolutionary conservation of larval eyes in Bilateria. Development 2002;129:1143–54.

[3] Ramirez MD, Pairett AN, Pankey MS, Serb JM, Speiser DI, Swafford AJ, et al. The last common ancestor of most bilaterian animals possessed at least nine opsins. Genome Biol Evol 2016;8:3640–52. doi:10.1093/gbe/evw248.

[4] Hardie RC, Juusola M. Phototransduction in *Drosophila*. Curr Opin Neurobiol 2015;34:37–45. doi:10.1016/j.conb.2015.01.008.

[5] Raible F, Tessmar-Raible K, Arboleda E, Kaller T, Bork P, Arendt D, et al. Opsins and clusters of sensory G-protein-coupled receptors in the sea urchin genome. Dev Biol 2006;300:461–75. doi:10.1016/j.ydbio.2006.08.070.

[6] Ullrich-Lüter EM, Dupont S, Arboleda E, Hausen H, Arnone MI. Unique system of photoreceptors in sea urchin tube feet 2011;108:8367–72. doi:10.1073/pnas.1018495108.

[7] Koyanagi M, Kubokawa K, Tsukamoto H, Shichida Y, Terakita A. Cephalochordate melanopsin: evolutionary linkage between invertebrate visual cells and vertebrate photosensitive retinal ganglion cells. Curr Biol 2005;15:1065–9. doi:10.1016/j.cub.2005.04.063.

[8] Sumner-Rooney L, Kirwan JD, Lowe E, Ullrich-Lüter E. Extraocular vision in a brittle star is mediated by chromatophore movement in response to ambient light. Curr Biol 2020;30:319–327.e4. doi:10.1016/j.cub.2019.11.042.

[9] Dzik J. Early Cambrian lobopodian sclerites and associated fossils from Kazakhstan. Palaeontology 2003;46:93–112. doi:10.1111/1475-4983.00289.

[10] Gehring WJ. Chance and necessity in eye evolution. Genome Biol Evol 2011;3:1053–66. doi:10.1093/gbe/evr061.

[11] Purschke G, Arendt D, Hausen H, Müller MCM. Photoreceptor cells and eyes in Annelida. Arthropod Struct Dev 2006;35:211–30. doi:10.1016/j.asd.2006.07.005.

[12] Couso JP. Segmentation, metamerism and the Cambrian explosion. Int J Dev Biol 2009;53:1305–16. doi:10.1387/ijdb.072425jc.

[13] Dray N, Tessmar-Raible K, Le Gouar M, Vibert L, Christodoulou F, Schipany K, et al. Hedgehog signaling regulates segment formation in the annelid Platynereis. 2010;329:339–42. doi:10.1126/science.1188913.

[14] Chen Z, Zhou C, Yuan X, Xiao S. Death march of a segmented and trilobate bilaterian elucidates early animal evolution. Nature 2019;573:412–5. doi:10.1038/s41586-019-1522-7.

[15] Senthilan PR, Piepenbrock D, Ovezmyradov G, Nadrowski B, Bechstedt S, Pauls S, et al. *Drosophila* auditory organ genes and genetic hearing defects. Cell 2012;150:1042–54. doi:10.1016/j.cell.2012.06.043.

[16] Zanini D, Giraldo D, Warren B, Katana R, Andrés M, Reddy S, et al. Proprioceptive Opsin functions in *Drosophila* larval locomotion. Neuron 2018;98:67–74.e4. doi:10.1016/j.neuron.2018.02.028.

[17] Backfisch B, Veedin Rajan VB, Fischer RM, Lohs C, Arboleda E, Tessmar-Raible K, et al. Stable transgenesis in the marine annelid *Platynereis dumerilii* sheds new light on photoreceptor evolution. Proc Natl Acad Sci USA 2013;110:193–8. doi:10.1073/pnas.1209657109.

[18] Baker GE, de Grip WJ, Turton M, Wagner H-J, Foster RG, Douglas RH. Light sensitivity in a vertebrate mechanoreceptor? J Exp Biol 2015;218:2826–9. doi:10.1242/jeb.125203.

[19] Fritzsch B, Piatigorsky J, Tessmar-Raible K, Jékely G, Guy K, Raible F, et al. Ancestry of photic and mechanic sensation? 2005;308:1113– 4. doi:10.1126/science.308.5725.1113.

[20] Bannister S, Antonova O, Polo A, Lohs C, Hallay N, Valinciute A, et al. TALENs mediate efficient and heritable mutation of endogenous genes in the marine annelid *Platynereis dumerilii*. Genetics 2014;197:77–89. doi:10.1534/genetics.113.161091.

[21] Gühmann M, Jia H, Randel N, Verasztó C, Bezares-Calderón LA, Michiels NK, et al. Spectral tuning of phototaxis by a Go-opsin in the rhabdomeric eyes of *Platynereis*. Curr Biol 2015;25:2265–71. doi:10.1016/j.cub.2015.07.017.

[22] Bezares-Calderón LA, Berger J, Jasek S, Verasztó C, Mendes S, Gühmann M, et al. Neural circuitry of a polycystin-mediated hydrodynamic startle response for predator avoidance. Elife 2018;7:e01668. doi:10.7554/eLife.36262.

[23] Fischer A, Dorresteijn A. The polychaete *Platynereis dumerilii* (Annelida): a laboratory animal with spiralian cleavage, lifelong segment proliferation and a mixed benthic/pelagic life cycle. Bioessays 2004;26:314–25.

[24] Randel N, Bezares-Calderón LA, Gühmann M, Shahidi R, Jékely G. Expression dynamics and protein localization of rhabdomeric opsins in *Platynereis* larvae. Integr Comp Biol 2013;53:7–16. doi:10.1093/icb/ict046.

[25] Picelli S, Faridani OR, Björklund AK, Winberg G, Sagasser S, Sandberg R. Full-length RNA-seq from single cells using Smart-seq2. Nat Protoc 2014;9:171–81. doi:10.1038/nprot.2014.006.

[26] Zantke J, Ishikawa-Fujiwara T, Arboleda E, Lohs C, Schipany K, Hallay N, et al. Circadian and circalunar clock interactions in a marine annelid. Cell Rep 2013;5:99–113. doi:10.1016/j.celrep.2013.08.031.

[27] Robinson MD, McCarthy DJ, Smyth GK. edgeR: a Bioconductor package for differential expression analysis of digital gene expression data. Bioinformatics 2010;26:139–40. doi:10.1093/bioinformatics/btp616.

[28] Tomer R, Denes AS, Tessmar-Raible K, Arendt D. Profiling by image registration reveals common origin of annelid mushroom bodies and vertebrate pallium. Cell 2010;142:800–9. doi:10.1016/j.cell.2010.07.043.

[29] Randel N, Asadulina A, Bezares-Calderón LA, Verasztó C, Williams EA, Conzelmann M, et al. Neuronal connectome of a sensory-motor circuit for visual navigation. Elife 2014;3:e02730. doi:10.7554/eLife.02730.

[30] Ayers T, Tsukamoto H, Gühmann M, Veedin Rajan VB, Tessmar-Raible K. A Go-type opsin mediates the shadow reflex in the annelid *Platynereis dumerilii*. BMC Biol 2018;16:41–9. doi:10.1186/s12915-018-0505-8.

[31] Fischer A, Brökelmann J. Das Auge von *Platynereis dumerilii* (Polychaeta): Sein Feinbau im ontogenetischen und adaptiven Wandel [The eye of *Platynereis dumerilii* (Polychaeta): Its fine structure in ontogenetic and adaptive change]. Z Zellforsch Mikrosk Anat 1966;71:217–44.

[32] Pende M, Vadiwala K, Schmidbaur H, Stockinger AW, Murawala P, Saghafi S, et al. A versatile depigmentation, clearing, and labeling method for exploring nervous system diversity. Science Advances 2020;6:eaba0365. doi:10.1126/sciadv.aba0365.

[33] Denes AS, Jékely G, Steinmetz PRH, Raible F, Snyman H, Prud’homme B, et al. Molecular architecture of annelid nerve cord supports common origin of nervous system centralization in Bilateria. Cell 2007;129:277–88. doi:10.1016/j.cell.2007.02.040.

[34] Kerner P, Simionato E, Le Gouar M, Vervoort M. Orthologs of key vertebrate neural genes are expressed during neurogenesis in the annelid *Platynereis dumerilii*. Evol Dev 2009;11:513–24. doi:10.1111/j.1525-142X.2009.00359.x.

[35] Yang Z, Edenberg HJ, Davis RL. Isolation of mRNA from specific tissues of *Drosophila* by mRNA tagging. Nucleic Acids Res 2005;33:e148–8. doi:10.1093/nar/gni149.

[36] Zou J, Zheng T, Ren C, Askew C, Liu X-P, Pan B, et al. Deletion of PDZD7 disrupts the Usher syndrome type 2 protein complex in cochlear hair cells and causes hearing loss in mice. Hum Mol Genet 2014;23:2374–90. doi:10.1093/hmg/ddt629.

[37] Boëda B, El-Amraoui A, Bahloul A, Goodyear R, Daviet L, Blanchard S, et al. Myosin VIIa, harmonin and cadherin 23, three Usher I gene products that cooperate to shape the sensory hair cell bundle. EMBO J 2002;21:6689–99. doi:10.1093/emboj/cdf689.

[38] Neef J, Jung S, Wong AB, Reuter K, Pangršič T, Chakrabarti R, et al. Modes and regulation of endocytic membrane retrieval in mouse auditory hair cells. J Neurosci 2014;34:705–16. doi:10.1523/JNEUROSCI.3313-13.2014.

[39] Stapelbroek JM, Peters TA, van Beurden DHA, Curfs JHAJ, Joosten A, Beynon AJ, et al. ATP8B1 is essential for maintaining normal hearing. Proc Natl Acad Sci USA 2009;106:9709–14. doi:10.1073/pnas.0807919106.

[40] Schneider ME, Dosé AC, Salles FT, Chang W, Erickson FL, Burnside B, et al. A new compartment at stereocilia tips defined by spatial and temporal patterns of myosin IIIa expression. J Neurosci 2006;26:10243–52. doi:10.1523/JNEUROSCI.2812-06.2006.

[41] Gou Y, Vemaraju S, Sweet EM, Kwon H-J, Riley BB. *Sox2* and *sox3* play unique roles in development of hair cells and neurons in the zebrafish inner ear. Dev Biol 2018;435:73–83. doi:10.1016/j.ydbio.2018.01.010.

[42] Kiernan AE, Cordes R, Kopan R, Gossler A, Gridley T. The Notch ligands DLL1 and JAG2 act synergistically to regulate hair cell development in the mammalian inner ear. Development 2005;132:4353–62. doi:10.1242/dev.02002.

[43] Oshima A, Suzuki S, Takumi Y, Hashizume K, Abe S, Usami S. *CRYM* mutations cause deafness through thyroid hormone binding properties in the fibrocytes of the cochlea. J Med Genet 2006;43:e25– 5. doi:10.1136/jmg.2005.034397.

[44] Tan J, Prakash MD, Kaiserman D, Bird PI. Absence of SERPINB6A causes sensorineural hearing loss with multiple histopathologies in the mouse inner ear. Am J Pathol 2013;183:49–59. doi:10.1016/j.ajpath.2013.03.009.

[45] Donaudy F, Snoeckx R, Pfister M, Zenner H-P, Blin N, Di Stazio M, et al. Nonmuscle myosin heavy-chain gene *MYH14* is expressed in cochlea and mutated in patients affected by autosomal dominant hearing impairment (DFNA4). Am J Hum Genet 2004;74:770–6. doi:10.1086/383285.

[46] Fu X, Zhang L, Jin Y, Sun X, Zhang A, Wen Z, et al. Loss of Myh14 increases susceptibility to noise-Induced hearing loss in CBA/CaJ Mice. Neural Plasticity 2016;2016:1–16. doi:10.1155/2016/6720420.

[47] Niwa N, Hiromi Y, Okabe M. A conserved developmental program for sensory organ formation in *Drosophila melanogaster*. Nat Genet 2004;36:293–7. doi:10.1038/ng1308.

[48] Fritzsch B, Beisel KW, Pauley S, Soukup G. Molecular evolution of the vertebrate mechanosensory cell and ear. Int J Dev Biol 2007;51:663– 78. doi:10.1387/ijdb.072367bf.

[49] Xiao T, Roeser T, Staub W, Baier H. A GFP-based genetic screen reveals mutations that disrupt the architecture of the zebrafish retinotectal projection. Development 2005;132:2955–67. doi:10.1242/dev.01861.

[50] Scott K, Becker A, Sun Y, Hardy R, Zuker C. Gq alpha protein function *in vivo*: genetic dissection of its role in photoreceptor cell physiology. Neuron 1995;15:919–27. doi:10.1016/0896-6273(95)90182-5.

[51] Bailes HJ, Lucas RJ. Human melanopsin forms a pigment maximally sensitive to blue light (λmax ≈ 479 nm) supporting activation of G(q/11) and G(i/o) signalling cascades. Proceedings of the Royal Society B: Biological Sciences 2013;280:20122987. doi:10.1098/rspb.2012.2987.

[52] Arendt D, Tessmar-Raible K, Snyman H, Dorresteijn AW, Wittbrodt J. Ciliary photoreceptors with a vertebrate-type opsin in an invertebrate brain. 2004;306:869–71. doi:10.1126/science.1099955.

[53] Bailes HJ, Zhuang L-Y, Lucas RJ. Reproducible and sustained regulation of Gαs signalling using a metazoan opsin as an optogenetic tool. PLoS One 2012;7:e30774. doi:10.1371/journal.pone.0030774.

[54] Govardovskii VI, Fyhrquist N, Reuter T, Kuzmin DG, Donner K. In search of the visual pigment template. Visual Neuroscience 2000;17:509–28. doi:10.1017/s0952523800174036.

[55] Street VA, McKee-Johnson JW, Fonseca RC, Tempel BL, Noben-Trauth K. Mutations in a plasma membrane Ca2+-ATPase gene cause deafness in deafwaddler mice. Nat Genet 1998;19:390–4. doi:10.1038/1284.

[56] Kozel PJ, Davis RR, Krieg EF, Shull GE, Erway LC. Deficiency in plasma membrane calcium ATPase isoform 2 increases susceptibility to noise-induced hearing loss in mice. Hear Res 2002;164:231–9. doi:10.1016/s0378-5955(01)00420-8.

[57] Klose MK, Boulianne GL, Robertson RM, Atwood HL. Role of ATP- dependent calcium regulation in modulation of *Drosophila* synaptic thermotolerance. J Neurophysiol 2009;102:901–13. doi:10.1152/jn.91209.2008.

[58] Chen X, Cuadros MD, Chalfie M. Identification of nonviable genes affecting touch sensitivity in *Caenorhabditis elegans* using neuronally enhanced feeding RNA interference. G3 (Bethesda) 2015;5:467–75. doi:10.1534/g3.114.015776.

[59] Arboleda E, Zurl M, Waldherr M, Tessmar-Raible K. Differential impacts of the head on *Platynereis dumerilii* peripheral circadian rhythms. Front Physiol 2019;10:900. doi:10.3389/fphys.2019.00900.

[60] Horridge GA. Proprioceptors, bristle receptors, efferent sensory impulses, neurofibrils and number of axons in the parapodial nerve of the polychaete *Harmothoë*. Proc R Soc Lond, B, Biol Sci 1963;157:199–222. doi:10.1098/rspb.1963.0005.

[61] Dorsett DA. The sensory and motor innervation of *Nereis*. Proceedings of the Royal Society of London B: Biological Sciences 1964;159:652–67. doi:10.1098/rspb.1964.0024.

[62] Schneider S, Fischer A, Dorresteijn AWC. A morphometric comparison of dissimilar early development in sibling species of *Platynereis* (*Annelida, Polychaeta*). Roux’s Arch Dev Biol 1992;201:243–56.

[63] Veedin Rajan VB, Häfker NS, Arboleda E, Poehn B, Gossenreiter T, Gerrard E, et al. Seasonal variation in UVA light drives hormonal and behavioral changes in a marine annelid via a ciliary opsin. Nat Ecol Evol in press. doi:10.1038/s41559-020-01356-1.

[64] Liang C, FANTOM Consortium, Forrest ARR, Wagner GP. The statistical geometry of transcriptome divergence in cell-type evolution and cancer. Nat Commun 2015;6:6066–6. doi:10.1038/ncomms7066.

[65] Arendt D, Musser JM, Baker CVH, Bergman A, Cepko C, Erwin DH, et al. The origin and evolution of cell types. Nat Rev Genet 2016;17:744– 57. doi:10.1038/nrg.2016.127.

[66] Arendt D, Bertucci PY, Achim K, Musser JM. Evolution of neuronal types and families. Curr Opin Neurobiol 2019;56:144–52. doi:10.1016/j.conb.2019.01.022.

[67] Piatigorsky J, Kozmik Z. Cubozoan jellyfish: an Evo/Devo model for eyes and other sensory systems. Int J Dev Biol 2004;48:719–29. doi:10.1387/ijdb.041851jp.

[68] Schlosser G. A short history of nearly every sense – The evolutionary history of vertebrate sensory cell types. Integr Comp Biol 2018;58:301–16. doi:10.1093/icb/icy024.

[69] Cosgrove D, Zallocchi M. Usher protein functions in hair cells and photoreceptors. The International Journal of Biochemistry & Cell Biology 2014;46:80–9. doi:10.1016/j.biocel.2013.11.001.

[70] Katana R, Guan C, Zanini D, Larsen ME, Giraldo D, Geurten BRH, et al. Chromophore-independent roles of Opsin apoproteins in *Drosophila* mechanoreceptors. Curr Biol 2019;29:2961–4. doi:10.1016/j.cub.2019.07.036.

[71] Hardie RC, Franze K. Photomechanical responses in Drosophila photoreceptors. 2012;338:260–3. doi:10.1126/science.1222376.

[72] Leung NY, Thakur DP, Gurav AS, Kim SH, Di Pizio A, Niv MY, et al. Functions of Opsins in *Drosophila* taste. Curr Biol 2020. doi:10.1016/j.cub.2020.01.068.

[73] Plachetzki DC, Fong CR, Oakley TH. The evolution of phototransduction from an ancestral cyclic nucleotide gated pathway. Proceedings of the Royal Society B: Biological Sciences 2010;277:1963–9. doi:10.1098/rspb.2009.1797.

[74] Plachetzki DC, Fong CR, Oakley TH. Cnidocyte discharge is regulated by light and opsin-mediated phototransduction. BMC Biol 2012;10:17– 10. doi:10.1186/1741-7007-10-17.

[75] Tessmar-Raible K, Steinmetz PR, Snyman H, Hassel M, Arendt D. Fluorescent two-color whole mount in situ hybridization in *Platynereis dumerilii* (Polychaeta, Annelida), an emerging marine molecular model for evolution and development. Biotechniques 2005;39:460–4.

[76] Hauenschild C, Fischer A. Platynereis dumerilii. Mikroskopische Anatomie, Fortpflanzung und Entwicklung [Platynereis dumerilii. Microscopical anatomy, reproduction and development]. Großes Zoologisches Praktikum, Stuttgart: G. Fischer Verlag; 1969.

[77] Revilla-i-Domingo R, Schmidt C, Zifko C, Raible F. Establishment of transgenesis in the demosponge *Suberites domuncula*. Genetics 2018;210:435–43. doi:10.1534/genetics.118.301121.

[78] Grabherr MG, Haas BJ, Yassour M, Levin JZ, Thompson DA, Amit I, et al. Full-length transcriptome assembly from RNA-Seq data without a reference genome. Nat Biotechnol 2011;29:644–52. doi:10.1038/nbt.1883.

[79] Sedlazeck FJ, Rescheneder P, Haeseler von A. NextGenMap: fast and accurate read mapping in highly polymorphic genomes. Bioinformatics 2013;29:2790–1. doi:10.1093/bioinformatics/btt468.

[80] Robinson MD, Oshlack A. A scaling normalization method for differential expression analysis of RNA-seq data. Genome Biol 2010;11. doi:10.1186/gb-2010-11-3-r25.

[81] Edgar RC. MUSCLE: multiple sequence alignment with high accuracy and high throughput. Nucleic Acids Res 2004;32:1792–7. doi:10.1093/nar/gkh340.

[82] Nguyen L-T, Schmidt HA, Haeseler von A, Minh BQ. IQ-TREE: a fast and effective stochastic algorithm for estimating maximum-likelihood phylogenies. Mol Biol Evol 2015;32:268–74. doi:10.1093/molbev/msu300.

[83] Inouye S, Noguchi M, Sakaki Y, Takagi Y, Miyata T, Iwanaga S, et al. Cloning and sequence analysis of cDNA for the luminescent protein aequorin. Proc Natl Acad Sci USA 1985;82:3154–8. doi:10.1073/pnas.82.10.3154.

[84] Virtanen P, Gommers R, Oliphant TE, Haberland M, Reddy T, Cournapeau D, et al. SciPy 1.0: fundamental algorithms for scientific computing in Python. Nat Methods 2020;17:261–72. doi:10.1038/s41592-019-0686-2.

[85] McKinney W. Data Structures for Statistical Computing in Python. SciPy, SciPy; 2010, pp. 56–61. doi:10.25080/Majora-92bf1922-00a.

[86] Harris CR, Millman KJ, van der Walt SJ, Gommers R, Virtanen P, Cournapeau D, et al. Array programming with NumPy. Nature 2020;585:357–62. doi:10.1038/s41586-020-2649-2.

[87] Doyle EL, Booher NJ, Standage DS, Voytas DF, Brendel VP, VanDyk JK, et al. TAL Effector-Nucleotide Targeter (TALE-NT) 2.0: tools for TAL effector design and target prediction. Nucleic Acids Res 2012;40:W117–22. doi:10.1093/nar/gks608.

[88] Cermak T, Doyle EL, Christian M, Wang L, Zhang Y, Schmidt C, et al. Efficient design and assembly of custom TALEN and other TAL effector-based constructs for DNA targeting. Nucleic Acids Res 2011;39:e82–2. doi:10.1093/nar/gkr218.

[89] Dahlem TJ, Hoshijima K, Jurynec MJ, Gunther D, Starker CG, Locke AS, et al. Simple methods for generating and detecting locus-specific mutations induced with TALENs in the zebrafish genome. PLoS Genet 2012;8:e1002861. doi:10.1371/journal.pgen.1002861.

